# Transdiagnostic electrophysiological subtypes reveal brain-behavior dimensions in youth psychiatry

**DOI:** 10.1101/2025.08.01.668189

**Authors:** Judie Tabbal, Aida Ebadi, Gabriel Robert, Borja Rodríguez-Herreros, Ahmad Mheich, Nadia Chabane, Aline Lefebvre, Anton Iftimovici, Sahar Allouch, Mahmoud Hassan

## Abstract

Conventional psychiatric diagnoses often fail to reflect the underlying neurobiological and behavioral complexity of mental health conditions. Here, we propose a transdiagnostic, data-driven framework for stratifying youth based on large-scale multisite electroencephalography (EEG) data from 1,707 individuals aged 5–18 years, including healthy controls and individuals diagnosed with attention-deficit/hyperactivity disorder (ADHD), autism spectrum disorder (ASD), anxiety disorders (ANX), and learning disabilities (LD), along with their common comorbidities. By applying normative modeling to quantify individual deviations from typical brain functional maturation, and integrating multidimensional EEG features across spectral, temporal, complexity, and dynamical domains via similarity network fusion clustering, we identified three robust neurophysiological biotypes. These biotypes showed distinct electrophysiological and behavioral profiles, and captured meaningful brain-behavior relationships. Our findings suggest that biologically informed subtypes capture meaningful neuropsychiatric heterogeneity in youth, challenging conventional diagnostic boundaries in psychiatric nosology.

## Introduction

Psychiatric diagnoses have traditionally been guided by symptom-based classification systems, notably the Diagnostic and Statistical Manual of Mental Disorders (DSM-5) and the International Classification of Diseases (ICD-11)^1^. While these taxonomies have provided a common language for clinical practice and research, they are increasingly criticized for their limited alignment with underlying neurobiological mechanisms^2–4^. Major concerns include high rates of comorbidity, considerable symptom overlap across diagnostic categories, and substantial heterogeneity within individuals with a specific neuropsychiatric diagnosis, which collectively call into question the validity of current nosological boundaries^3^. Furthermore, findings from neuroimaging and electrophysiological studies suggest that brain-based biomarkers often fail to align with existing diagnostic categories^2,5,6^. This misalignment poses a significant barrier to advancing our understanding of the neurobiological substrates of mental illness and hampers the development of targeted treatments. In response, biologically grounded dimensional frameworks such as the Research Domain Criteria (RDoC)^3^ have generated substantial interest over the past decade. Yet, their anticipated clinical utility has largely fallen short.

Against this backdrop, considerable efforts have been devoted to developing data-driven, biologically informed methodologies capable of capturing the transdiagnostic complexity of mental disorders. However, most traditional analyses still rely on group-level comparisons, which assume homogeneity within diagnostic categories. Such an approach obscures important inter-individual variability and may overlook subtle but clinically relevant patterns. Averaging data across diagnostic groups can dilute or mask individual-specific biomarkers, thereby limiting the clinical applicability of the findings. Normative modeling (NM) has recently been introduced in neuroimaging as a promising alternative, offering a framework to quantify individual deviations from population-derived normative distributions. Unlike conventional case-control designs, NM provides subject-level deviation scores enabling the detection of neurophysiological alterations at the individual level^7,8^. NM has demonstrated its utility in magnetic resonance imaging (MRI) for mapping lifespan trajectories of grey and white matter volume, cortical thickness, and surface area^8–10^, as well as for characterizing structural heterogeneity in autism, schizophrenia, ADHD, bipolar disorder, and Alzheimer’s disease^11–14^. Efforts have also begun to apply NM to functional MRI and EEG data, albeit to a lesser extent^8,15,16^. Our recent work^17,18^ represents one of the first attempts to apply this approach to EEG data in psychiatric and neurodegenerative populations. In the present study, we build upon this foundation by leveraging individual-level deviation profiles to identify biologically grounded subtypes that transcend conventional diagnostic boundaries.

Using a harmonized, multisite high-density EEG (HD-EEG) dataset comprising 1,701 participants, including healthy controls (HC) and individuals diagnosed with attention-deficit/hyperactivity disorder (ADHD), autism spectrum disorder (ASD), anxiety (ANX), and learning disorders (LD), as well as their comorbidities, we computed individualized deviation scores across a comprehensive set of EEG features. The resulting multivariate deviation profiles served as input to similarity network fusion (SNF) clustering^19^, enabling the identification of subgroups with unique and distinctive electrophysiological patterns. To assess the clinical relevance of these subtypes, we analyzed their diagnostic profiles, comorbidity patterns, and demographic distributions. Additionally, we examined behavioral differences between clusters using dimensional phenotypic measures covering domains such as attention, depression, cognitive performance, and language abilities. We further investigated the contribution of individual EEG features to cluster formation and evaluated their predictive value for behavioral traits within each subgroup. This integrative, transdiagnostic approach enables the identification of biologically grounded subtypes and sheds light on their specific neurophysiological and behavioral characteristics.

## Results

### Large-scale, multisite EEG data and multimodal feature extraction provide a robust foundation for transdiagnostic analysis

High-density resting-state, eyes-open EEG data were collected from 1,701 subjects (44% M) aged 5 to 18 years, across five independent datasets (see “Methods”). Normative models were trained on 447 healthy control participants (HC_tr), with an independent set of 111 healthy controls (HC_te) held out as a test group for comparison against the clinical cohort. The clinical groups included 1,143 patients diagnosed with different psychiatric conditions, including ADHD, ASD, ANX, and LD. Many patients exhibited comorbidities involving two or more of these disorders. An overview of the sample distribution by age, sex, diagnostic group, site, and comorbidity status is provided in Fig. 1. Further details on data sample distribution across sites, as well as corresponding medication status and selection criteria, can be found in Supplementary Data 1.

**Fig. 1:**
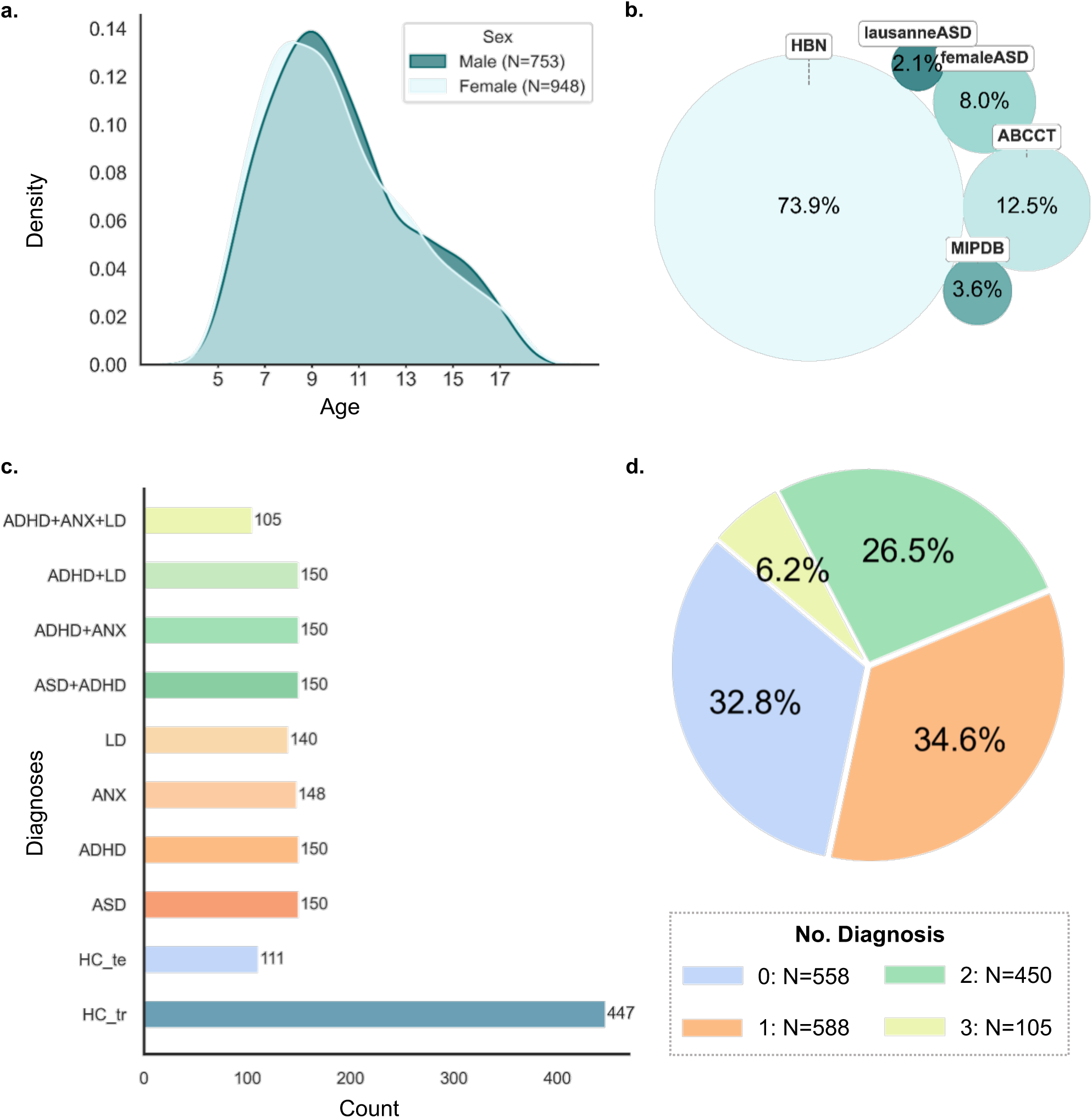
Overview of sample characteristics. **a** Age and sex distribution. **b** Distribution of participants across acquisition sites. **c** Diagnostic group composition. **d** Comorbidity profiles among participants.

After preprocessing and downsampling the EEG data to the standard 10-20 montage (19 channels), we extracted a comprehensive set of features, yielding a total of 1957 features, corresponding to 103 features per channel. These features were grouped into four main categories: (1) spectral features (e.g., band-specific power, aperiodic 1/f slope and offset, power ratios), (2) complexity features (e.g., entropy measures, Hurst exponent, detrended fluctuation analysis), (3) time-domain features (e.g., skewness, kurtosis, mean envelope, amplitude-based power), and (4) catch24 (a subset of time-series features from the ‘*hctsa’* toolbox^20,21^ (Fig. 2a-b). Details on the specific features within each category are outlined in Supplementary Data 2. Given that the data were acquired from multiple independent sites, site-related variability was addressed through feature harmonization using NeuroCombat^22^. Fig. 2c illustrates the distribution of selected features (mean across channels; one representative feature per category) before and after harmonization. Prior to harmonization, 101 out of 103 features (98%) exhibited significant inter-site variability (Kruskal-Wallis, *p* < 0.05), whereas after site harmonization, only 14 features (13%) remained significantly different across sites. This indicates that approximately 85% of the features were successfully adjusted for site effects.

**Fig. 2:**
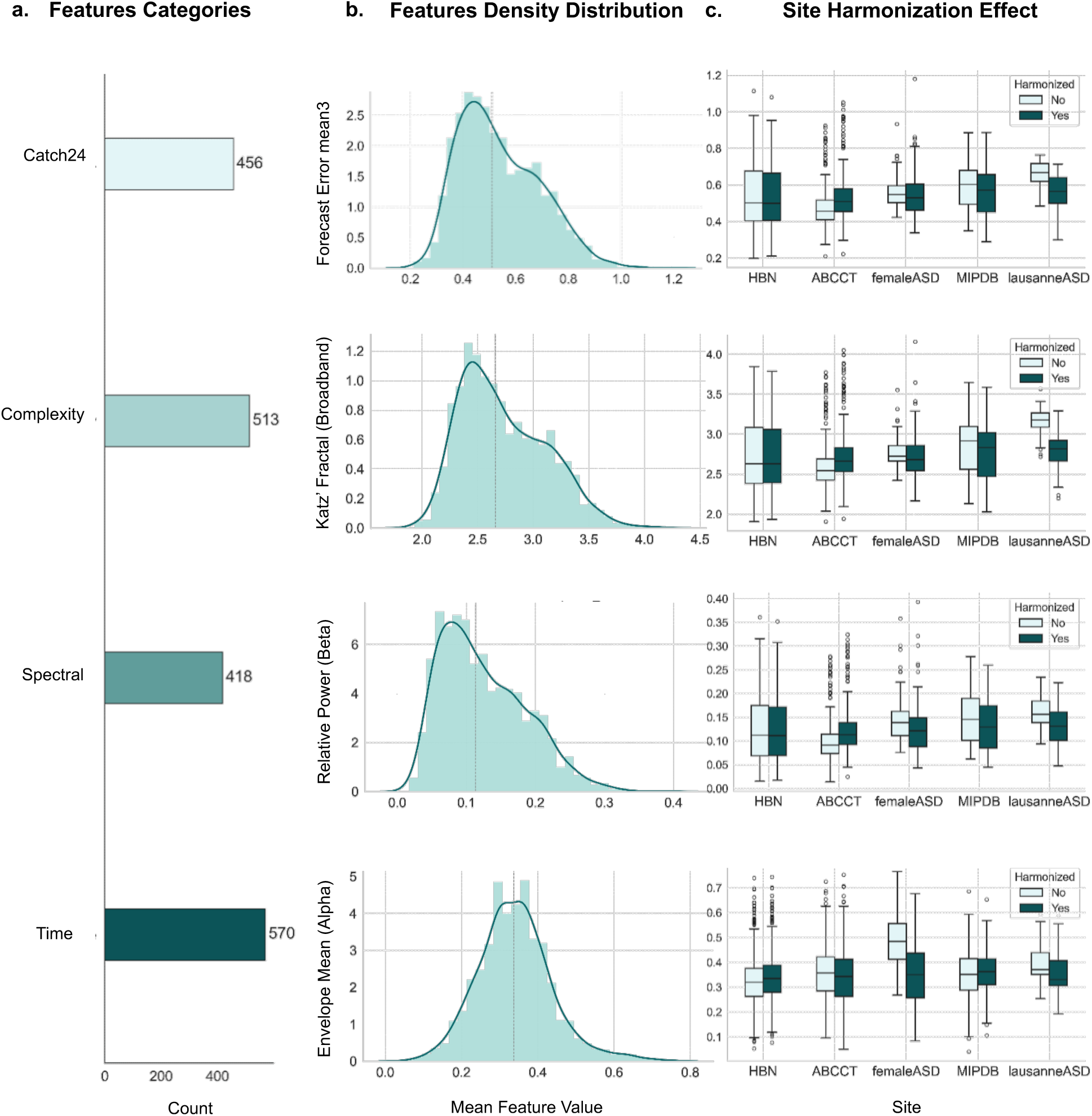
Overview of EEG features distribution and site harmonization. **a** Total number of extracted features per category (Catch24, complexity, spectral, and time). **b** Density plots of representative features (averaged across channels) for each category. **c** Distributions of the same features shown in b, before (light green) and after (dark green) site harmonization.

### Normative modeling reveals individualized deviations from typical brain function trajectory across healthy and clinical populations

We trained a Generalized Additive Model for Location, Scale, and Shape (GAMLSS) for each EEG feature using data from a reference cohort of 447 healthy controls (HC) (see “Methods”).

This allowed us to model normative electrophysiological trajectories spanning the age range of 5 to 18 years. Representative trajectories are shown in Fig. 3a, with the median and the 5th and 95th percentiles delineating the expected range of variation within the healthy population. Supplementary Data 3 provides details on the optimal distribution families, model parameters, and covariates used for each feature. Normal quantile-quantile (QQ) plots (Fig. 3c) indicate a good model fit, with residuals closely approximating a normal distribution. After model fitting, we projected all individuals from the clinical cohort, along with a held-out HC test set, onto the normative models to compute subject-specific deviation scores (z-scores). This yielded a matrix of standardized z-scores across all EEG features, channels, and subjects.

**Fig. 3:**
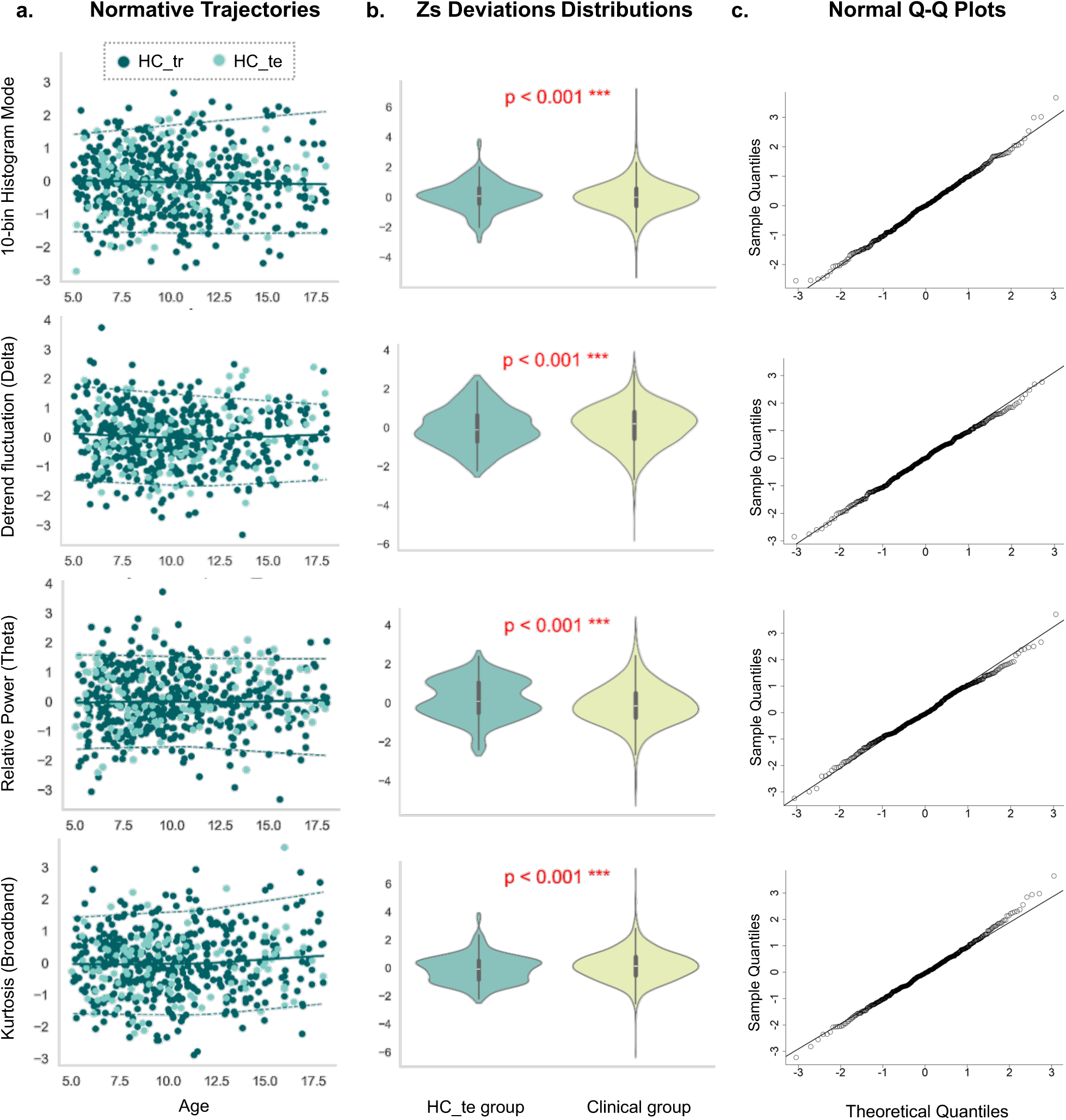
Overview of the normative modeling framework. **a** Normative trajectories of representative EEG features, derived from healthy controls in the training set (HC_tr), and tested on an independent testing set (HC_te), spanning ages 5–18 years. **b** Distribution of z-scores in the HC_te group compared to the clinical group. To ensure balanced sample size, bootstrapped undersampling was applied; red asterisks indicate statistically significant group differences. **c** Normal quantile-quantile plots comparing the sample quantiles to the theoretical normal distribution. Alignment along the diagonal line (slope = 1) indicates good model fit and normal distribution of deviations.

Figure 3b displays the distribution of z-scores for selected EEG features in the HC and clinical groups. In the HC test group, z-scores were largely confined within the normative range (±2), whereas the clinical group showed more pronounced deviations. Statistical comparisons revealed significant group differences for 89 out of 103 features (∼86.41%; Mann–Whitney U test, bonferroni-corrected p < 0.05), reflecting widespread atypicality in the clinical population. To mitigate sample size imbalance, we employed a bootstrapping strategy involving 100 iterations of random undersampling of the clinical cohort.

### Transdiagnostic clustering identifies distinct neurophysiological subtypes across healthy and clinical populations

To identify transdiagnostic neurophysiological subtypes, we applied unsupervised clustering to the EEG-based deviation scores using Similarity Network Fusion (SNF), which integrates multi-dimensional feature spaces into a unified similarity network, followed by spectral clustering (see “Methods”). The eigen-gap and rotation cost criteria, indicated local optima at *k*=2 and *k*=3 clusters (Fig. 4a). While a two-cluster solution is common in prior SNF applications ^23,24^, it often results in coarse groupings that may obscure finer-grained structure. We selected k=3 as the optimal solution, yielding three neurophysiological subtypes: Cluster C1 (n=581, 46.3%), Cluster C2 (n=400, 31.9%), and Cluster C3 (n=273, 21.8%). Visualization of the fused similarity matrix showed strong intra-cluster cohesion and clear inter-cluster separation (Fig. 4c). Notably, cluster C3 exhibited the highest internal homogeneity, and moderate overlap was observed between clusters C2 and C3. The affinity matrices derived from individual feature categories (Supplementary Fig. S2) displayed convergent structure.

**Fig. 4:**
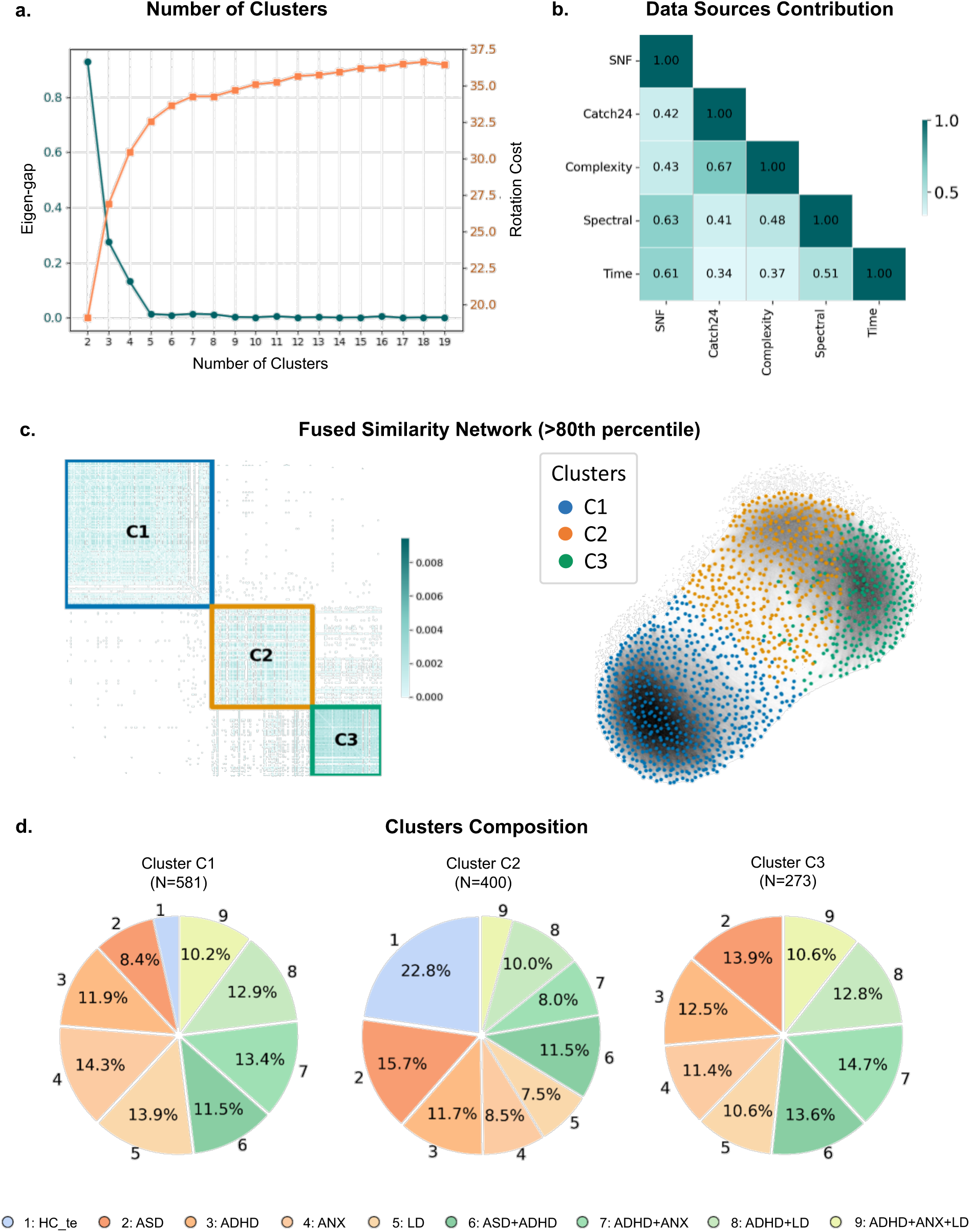
Transdiagnostic clustering results using Similarity Network Fusion (SNF). **a** Selection of the optimal number of clusters based on eigengap and rotation cost metrics. **b** Heatmap of Normalized Mutual Information (NMI) scores across feature categories, where higher values indicate higher contribution to the fused similarity network. **c** Fused similarity network visualized as a heatmap and a graph, thresholded at the 80th percentile of similarity strength. Nodes represent participants, and edges represent pairwise similarity strength. **d** Proportions of diagnostic groups within each cluster.

All EEG feature categories contributed comparably to the fused similarity network. Spectral and time-domain features exhibited slightly greater influence, as indicated by Normalized Mutual Information (NMI) scores of 0.63 and 0.61, respectively. Complexity and Catch24 features contributed to a lesser extent, with NMI values of 0.43 and 0.41, respectively (Fig. 4b). At the individual feature level, contributions were further assessed, and the most informative features are presented in Supplementary Fig. S6.

To assess the clinical relevance of the identified clusters, we examined their diagnostic composition (Fig. 4d). Cluster C1 was characterized by individuals with ANX (14.3%) and LD (13.9%), along with notable proportions of comorbid cases with ADHD, including ADHD+ANX (13.4%) and ADHD+LD (12.9%). Cluster C2 was predominantly composed of HC (22.8%), followed by individuals with ASD (15.7%). In contrast, cluster C3 was largely dominated by individuals with ADHD and its comorbidities, including ADHD+ANX (14.7%), ASD+ADHD (13.6%), ADHD+LD (12.8%), and ADHD alone (12.5%), along with a substantial proportion of ASD (13.9%). Notably, cluster C3 did not include any HC individuals. A Chi-square test of independence confirmed that the distribution of diagnostic categories across clusters was statistically significant (χ² = 180.67, df = 16, p < 0.0001), indicating that the observed cluster compositions are unlikely to have occurred by chance.

The identified clusters did not exhibit significant associations with key demographic variables. As shown in Supplementary Table S1, there were no statistically significant differences or moderate-to-large effect sizes across clusters with respect to age, sex, acquisition site, or comorbidity, suggesting that the clustering solution was not affected by these variables. Covariate distributions across clusters are further visualized in Supplementary Fig. S3. Moreover, the SNF clustering demonstrated high robustness to the subsampling of the original population, with an average cluster assignment stability of 97.62% across 100 iterations (see “Methods” for details).

### Neurophysiological subtypes exhibit unique electrophysiological signatures

To elucidate the electrophysiological underpinnings of the derived subtypes, we examined the distribution of deviation scores for the top EEG features, averaged across channels, that contributed most to the clustering, as ranked by their NMI scores. The top 15 features (Fig. 5a) explain approximately 25% of the variance relevant to clustering, while the top 50 features (Supplementary Fig. S4) account for around 70%. Notably, the top-ranked features included the longest stretch of decreasing values, measuring signal monotonic stability and temporal regularity, followed by entropy measures (approximate and sample entropy) indicating signal complexity and unpredictability, and adjusted gamma power representing high-frequency neural oscillations linked to cognitive processing.

**Fig. 5:**
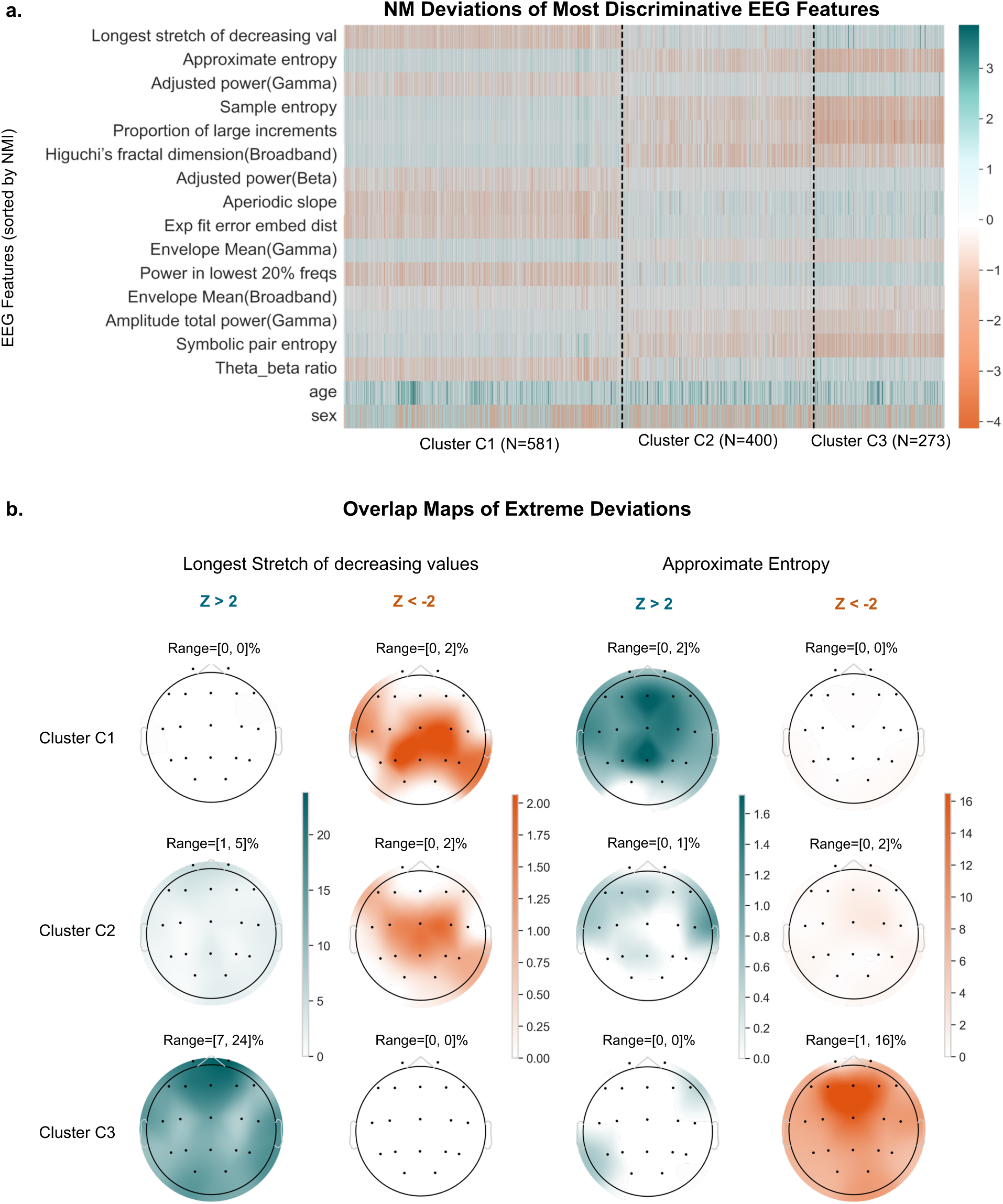
Cluster-specific electrophysiological signatures. **a** Heatmap representing the z-scores of the top 15 EEG features (averaged across channels) sorted by NMI scores (top to bottom) across clusters. Green indicates positive deviations and orange indicates negative deviations relative to normative models. **b** Scalp topographies showing the percentage of subjects with extreme deviations (Z>2: Green; Z<-2: orange) across the 19 EEG channels for the two most contributing features.

Distinct deviation patterns emerge across clusters: clusters C1 and C3 exhibit more pronounced and often oppositely directed deviations, whereas cluster C2 demonstrates distributions more closely aligned with normative values. Interestingly, the directionality of deviations in C2 generally mirrors that observed in C3. The distributions of age and sex across clusters are shown alongside, indicating a balanced representation and ruling out confounding by these demographic variables (Supplementary Table. S1). To spatially localize these electrophysiological deviations, we generated topographical overlap maps for the most discriminative EEG features (Fig. 5b). These maps illustrate the proportion of individuals within each cluster exhibiting extreme deviations (positive: z > 2; negative: z < −2) across the standard 19-channel EEG montage. For instance, the longest stretch of decreasing values showed a high prevalence of positive extreme deviations in cluster C3 (ranging from 7% to 24%) across the scalp, whereas deviation prevalence was negligible in C2 and absent in C1. Likewise, approximate entropy revealed negative extreme deviations in up to 16% of individuals within cluster C3, particularly concentrated over fronto-central electrodes.

### Identified subtypes align with specific behavioral profiles

We next investigated whether the neurophysiological subtypes identified via clustering of EEG-based deviation scores were associated with distinct behavioral profiles. Behavioral scores across 13 domains were compared among the three clusters. Significant differences were observed in 10 out of 13 domains using the Kruskal–Wallis H test (p < 0.05), including Hyperactivity, Attention, Depression, Internalizing and Externalizing symptoms, Anxiety, Social Functioning, Autistic Traits, Executive Function, Cognitive Function, and Emotional Reactivity (see Supplementary Table S3). Post hoc Dunn’s tests further revealed specific pairwise differences between clusters for several of these domains (Supplementary Fig. S5), indicating that the identified subtypes are behaviorally distinct and clinically relevant.

To illustrate inter-cluster behavioral differences, Fig. 6a displays the distribution of composite z-scores for the behavioral domains that significantly differed across clusters. Distinct profiles emerged: Cluster C1, comprising a higher proportion of individuals with anxiety and learning disorders, was marked by pronounced deficits in executive function and elevated emotional reactivity. Cluster C3, which included a substantial number of individuals with ADHD and common comorbidities, exhibited notably higher scores in hyperactivity-related domains. In contrast, Cluster C2, predominantly composed of healthy controls, showed the lowest composite z-scores across multiple domains, including attention, depression, internalization, anxiety, and social functioning, reflecting a broadly normative behavioral profile.

**Fig. 6:**
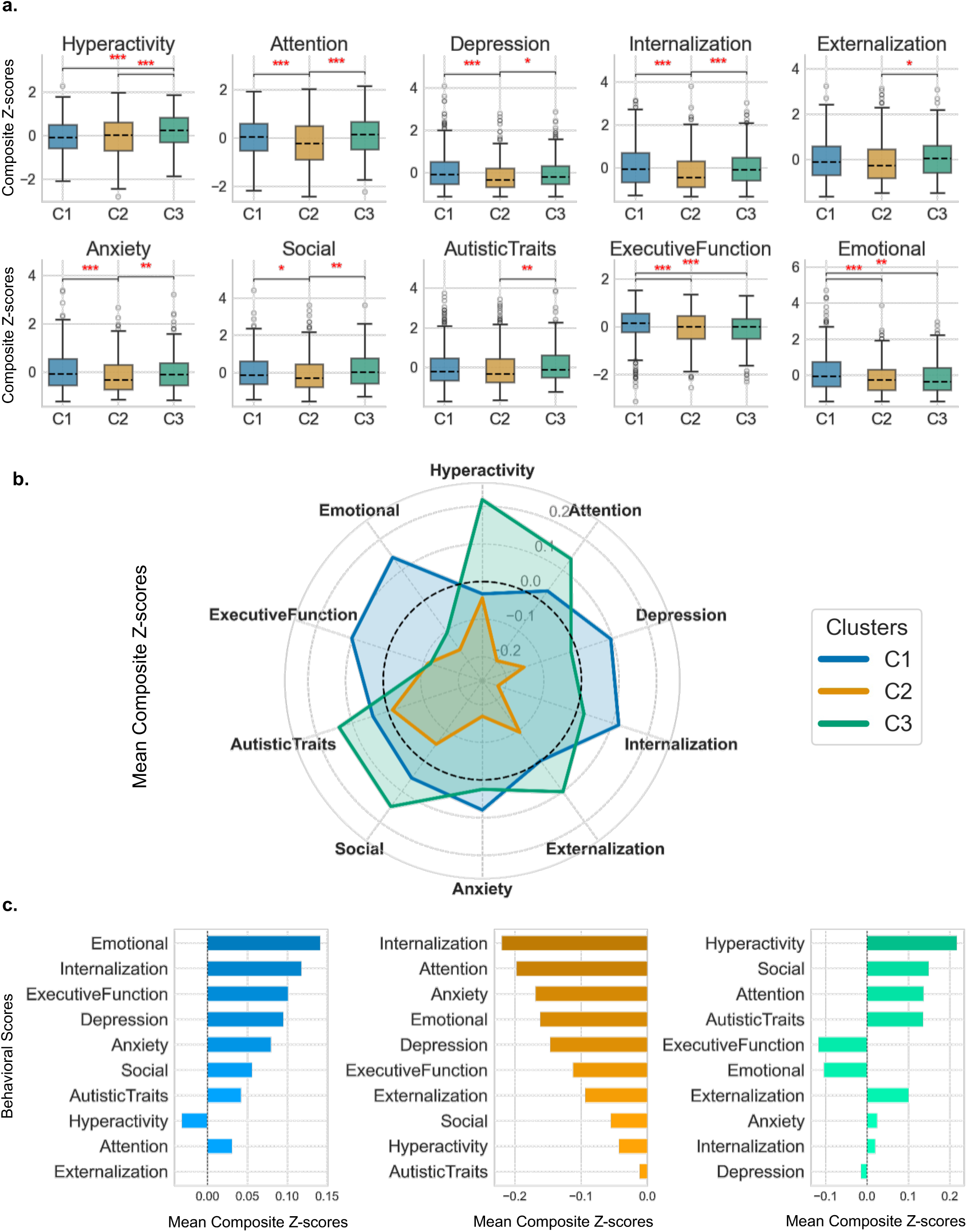
Cluster-specific behavioral profile. **a** Boxplots showing the distribution of individuals’ composite standardized scores for each behavioral domain across the three clusters. Red asterisks denote statistically significant differences between clusters (*p<0.05, **p<0.01, ***p<0.001). **b** Radar plot of mean composite standardized scores across behavioral domain for each cluster. The dashed dark circle indicates the reference value of zero; positive values reflect greater severity of behavioral problems relative to the average, and negative values reflect lower severity. **c** Bar plots depicting mean composite standardized scores for each behavioral domain, separately for each cluster.

To summarize the behavioral profile of each neurophysiological subtype, we computed the mean composite z-score for each behavioral domain within each cluster and ranked the domains by magnitude (Fig. 6b,c). Cluster C1 was predominantly characterized by elevated scores in emotional reactivity, internalizing symptoms, and executive dysfunction. Cluster C3 exhibited a distinct pattern of increased hyperactivity and social-attentional difficulties. In contrast, Cluster C2 displayed generally low scores across all behavioral domains, representing a reference or normative behavioral profile within the sample. To further assess the discriminative power of each behavioral domain, we calculated the proportion of variance explained across clusters. Internalization accounted for the largest share of explained variance, followed by attention, emotional dysregulation, and hyperactivity (Supplementary Fig. S6), identifying the behavioral dimensions most strongly associated with neurophysiological subtype differentiation.

### EEG-derived features predict cluster-specific clinical and behavioral traits

To further investigate the relationship between neurophysiological deviations and behavioral phenotypes, we conducted multiple linear regression (MLR) analyses within each cluster, focusing on the two most representative behavioral domains per subtype. All models showed statistically significant fits (p < 0.05), with correlation coefficients of r = 0.53 and r = 0.51 for emotional dysregulation and internalization in cluster C1, r = 0.62 and r = 0.75 for internalization and attention in cluster C2, and r = 0.75 for both hyperactivity and social functioning in cluster C3 (Fig. 7). Regression coefficients for the top 15 EEG features contributing to each behavioral domain are visualized in Fig. 7, ranked by absolute magnitude to indicate relative importance. Complete ranked lists of the top 50 features per domain are provided in Supplementary Fig. S7– S9. These findings highlight specific EEG features that significantly predict behavioral variability within subtypes, supporting a direct link between electrophysiological deviations and domain-specific symptomatology.

**Fig. 7:**
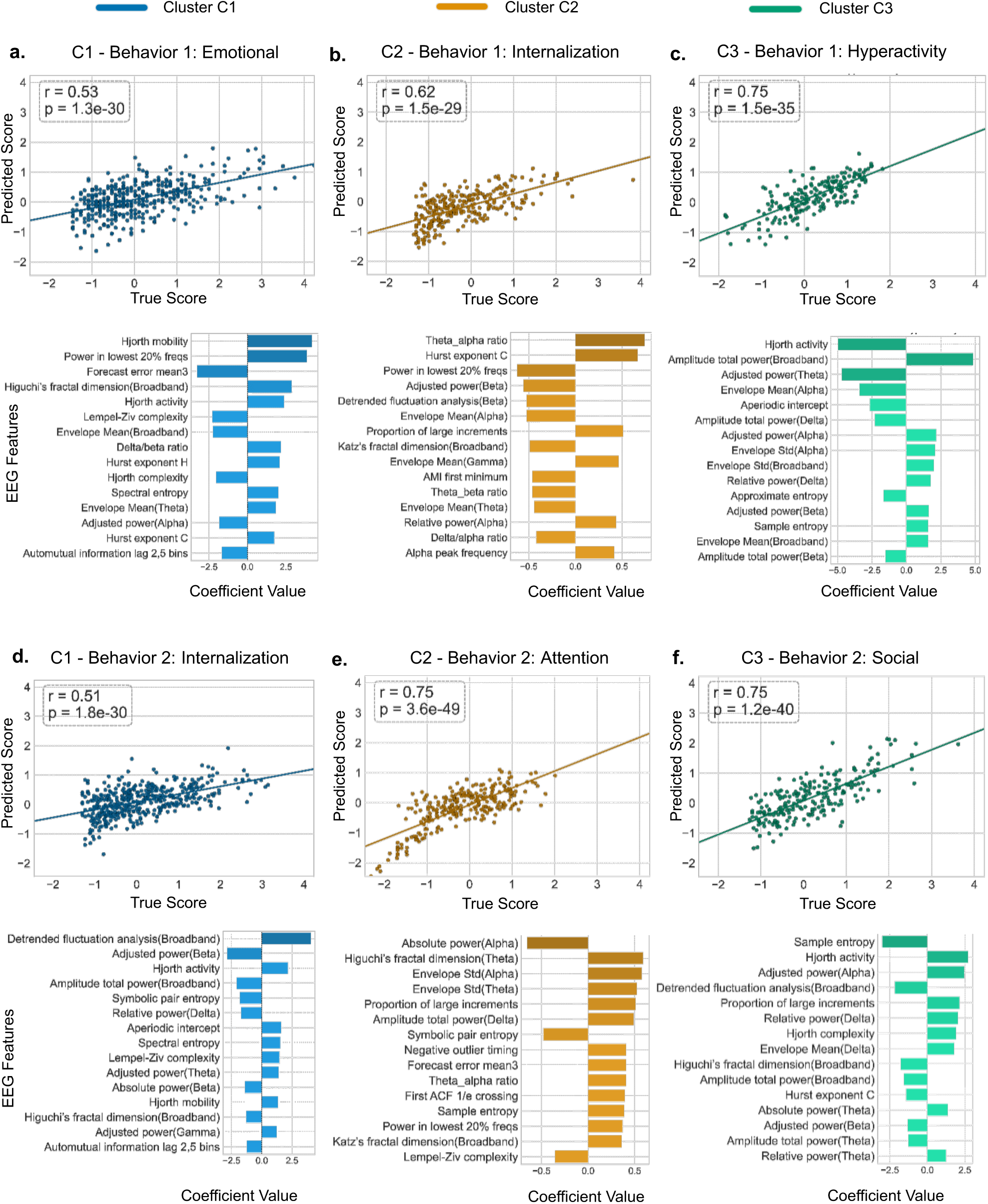
Multiple Linear Regression (MLR) analysis results for the top two prominent behavioral traits per cluster. Regression scatter plots show the predicted behavioral score resulted from the EEG features against/versus the true behaviroal score. Associated barplots illustrate standardized regression coefficients for the top 15 EEG features contributing to the prediction, ranked from highest (top) to lowest (bottom) absolute values.

**Fig. 8:**
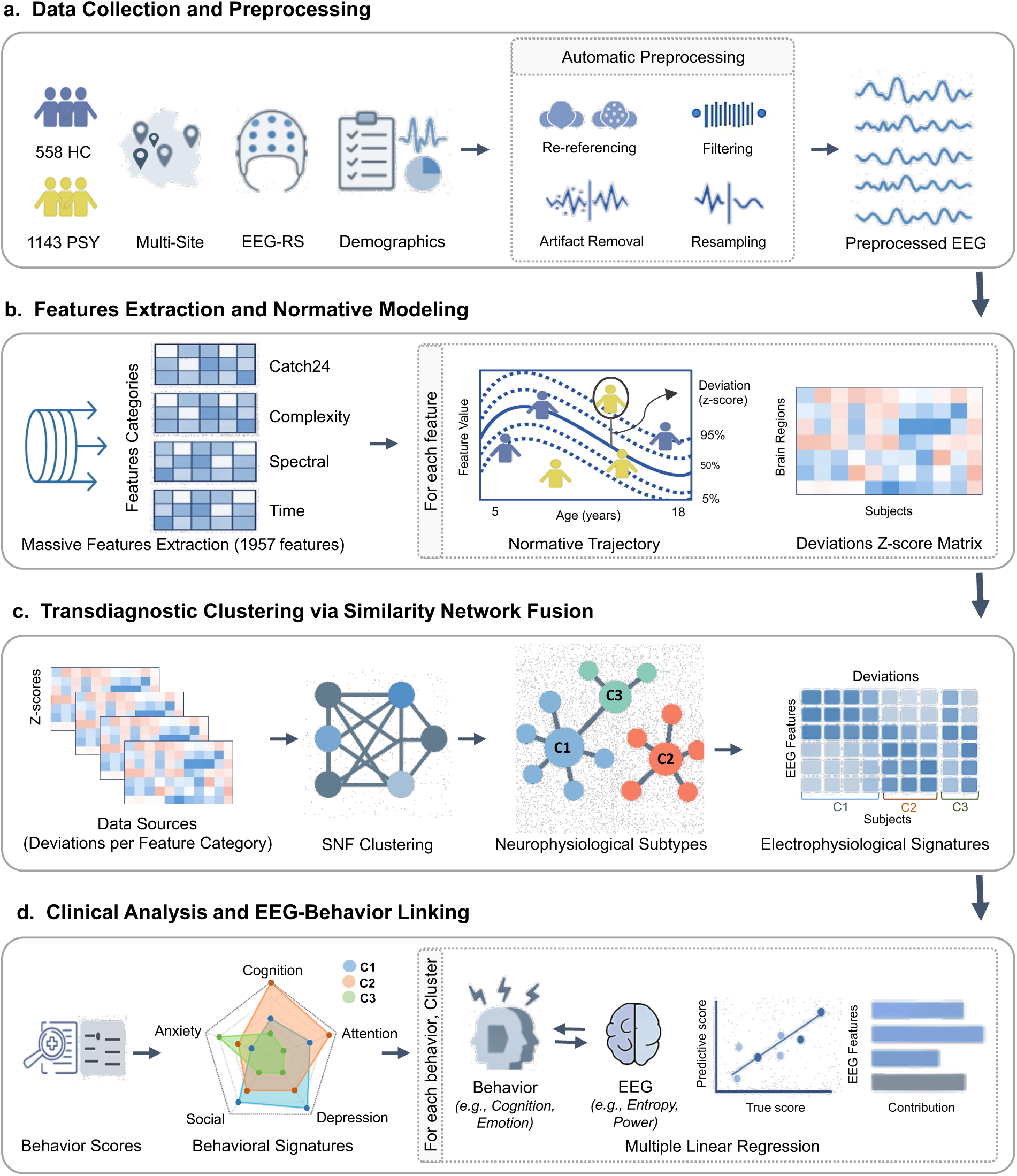
Overview of the study pipeline involving the main analysis steps. HC, Healthy Controls; PSY, Psychiatry; EEG-RS, Electroencephalography-RestingState; SNF, Similarity Network Fusion; C1, Cluster 1; C2, Cluster 2; C3, Cluster 3.

## Discussion

Current psychiatric diagnostic systems are primarily symptom-based and often overlook the underlying neurophysiological heterogeneity characteristic of neurodevelopmental disorders. In this study, we introduce a transdiagnostic, data-driven framework that integrates large-scale, multisite high-density EEG data. By applying normative modeling to quantify individualized deviations from typical neurodevelopmental trajectories, and using similarity network fusion to cluster multidimensional EEG features, we identified three robust and clinically meaningful neurophysiological subtypes spanning both healthy individuals and those with psychiatric diagnoses. These subtypes demonstrated distinct electrophysiological signatures, behavioral phenotypes, and brain-behavior associations, providing a biologically informed framework to parse diagnostic overlap and heterogeneity in neurodevelopmental psychiatric conditions.

A key finding of this study is that the EEG-derived subtypes identified through data-driven clustering did not align neatly with traditional diagnostic categories. Instead, each cluster comprised individuals from multiple diagnostic groups, including comorbid cases, underscoring the presence of shared neurophysiological characteristics across conventional diagnostic boundaries. This transdiagnostic overlap supports dimensional models such as the RDoC^5^, which emphasize underlying constructs over categorical diagnoses. Our findings are consistent with growing evidence that data-driven methods can uncover biologically meaningful subgroups that transcend standard diagnostic labels. For example, cross-diagnostic neuroimaging subtypes have been reported for internalizing symptoms in youth^25^, longitudinal trajectory-based clusters in early psychosis and depression^26^, and SNF-based subtypes spanning schizophrenia, bipolar disorder, autism spectrum disorder, and healthy controls^27,28^. These findings are reinforced by recent methodological reviews advocating for computational approaches to address psychiatric heterogeneity and support brain-based transdiagnostic stratification^29–31^, as well as by calls for biologically informed subtyping to enable precision psychiatry and targeted interventions^32^.

The distinct electrophysiological signatures associated with each subtype further highlight their biological differentiation. Cluster C2, encompassing the majority of healthy controls, exhibited deviation scores closely aligned with normative EEG trajectories, supporting its characterization as a neurotypical reference group. In contrast, clusters C1 and C3—enriched for individuals with internalizing symptoms (e.g., anxiety) and externalizing diagnoses (e.g., ADHD), respectively— displayed more pronounced and often opposing deviations across key EEG features. This divergence is consistent with prior literature suggesting antagonistic neurodevelopmental profiles between internalizing and externalizing disorders, despite overlapping symptomatology^33^. For example, features such as gamma band total power, envelope mean, and entropy measures were markedly elevated in cluster C3 but reduced in C1. These findings align with prior work implicating aberrant gamma activity in disrupted cortical excitation, impaired cognitive control, and altered network integration in ADHD^34,35^. Additionally, entropy-based features correspond with emerging evidence linking EEG complexity to emotional dysregulation^36^, cognitive rigidity (Keshmiri, 2020), and impulsivity across psychiatric populations^37,38^, supporting their potential as transdiagnostic biomarkers of psychopathology. Subtype-specific differences were also evident in spectral parameters such as adjusted gamma power, the aperiodic (1/f) component, and the theta/beta ratio. Alterations in the aperiodic slope have been linked to disrupted excitation/inhibition balance and inefficient neural processing in ADHD^39^, while the theta/beta ratio has shown prognostic utility in ADHD subgroups^40^. Although direct empirical associations between Catch24 features and psychiatric constructs are limited, the most discriminative among them—capturing signal stability, temporal regularity, and dynamic embedding—echo findings that suggest disruptions in these temporal characteristics may reflect altered functional connectivity across neuropsychiatric conditions^41^, warranting further investigation into their translational relevance.

Beyond their neural profiles, the identified subtypes demonstrated distinct behavioral phenotypes, underscoring their clinical relevance. C1 showed elevated emotional dysregulation and internalizing symptoms, reflecting core features of anxiety disorders and their strong association with emotional-internalizing psychopathology in youth^42–44^. Additionally, the high prevalence of learning disorders aligns with the dominance of executive function deficits in this cluster, consistent with evidence identifying executive dysfunction as central to learning disabilities^45^. C3 exhibited marked impairments in hyperactivity, social, and attention, mirroring behavioral profiles commonly observed in ADHD and ASD populations^46–49^. Conversely, C2, largely composed of typically developing individuals, showed minimal behavioral symptoms across domains. Importantly, multivariate models revealed that core behavioral traits per cluster were significantly predicted by distinct EEG features. These results highlight subtype-specific brain–behavior coupling mechanisms and suggest the clinical relevance of EEG-informed stratification.

Notably, the most influential EEG features distinguishing subtypes and predicting core behavioral traits, though distinct, spanned all major EEG domains—including Catch24, complexity, spectral, and time-domain measures—underscoring the value of multidimensional EEG representations in capturing the electrophysiological heterogeneity underlying psychiatric conditions.

Although our framework showed promising results with clinical relevance, several limitations warrant consideration. First, the original clinical cohort was considerably imbalanced across diagnostic groups, leading to the exclusion of a substantial number of samples during resampling—particularly from ADHD and multi-diagnoses groups—which reduced both sample size and comorbidity diversity. Second, validation of the entire analytical pipeline on independent external cohorts is necessary to confirm its reproducibility and reliability. Third, the cross-sectional nature of the data limits our ability to explore longitudinal developmental trajectories and track the dynamic evolution of neurophysiological subtypes over time. Moreover, future studies are encouraged to incorporate additional covariates, such as genetic, environmental, and behavioral factors, to enhance the explanatory power and interpretability of the models. Finally, the generalizability of these findings to the adult population remains unclear and should be explored in subsequent investigations.

Together, our findings demonstrate that EEG-based, data-driven stratification can identify biologically grounded subtypes that transcend traditional diagnostic categories, offering a principled framework to advance transdiagnostic research in psychiatry.

## Methods

The full analysis pipeline is illustrated in Fig. 8. It includes data collection and preprocessing, feature extraction, normative modeling, and subtype identification, followed by behavioral validation. Each step is described in detail in the following sections.

### Participants

Our study comprised 1,701 children and adolescents (ages 5–18 years; mean = 10.14 ± 3.03; 44% male), drawn from five large-scale, independent studies: the Healthy Brain Network (HBN)^50,51^, the Multimodal Resource for Studying Information Processing in the Developing Brain (MIPDB)^51^, the Autism Biomarkers Consortium for Clinical Trials (ABCCT)^52^, the Multimodal Developmental Neurogenetics of Females with ASD (femaleASD)^53^, and the LausanneASD study^54^. The initial dataset included 558 healthy controls (HC) and 2,316 patients clinically diagnosed with one or more psychiatric conditions including attention-deficit/hyperactivity disorder (ADHD), autism spectrum disorder (ASD), anxiety disorders (ANX), and learning disabilities (LD). Given the high prevalence of comorbid diagnoses (950 out of 2,316 patients; 41% presenting with more than one diagnosis), we considered each diagnosis, whether occurring alone or in combination with others, as a distinct clinical subgroup (e.g., ADHD, ASD+ADHD, ADHD+LD). To mitigate severe diagnostic class imbalance and ensure analytical robustness, we applied a structured resampling procedure to the patient cohort. Diagnostic subgroups with more than 150 participants were randomly downsampled to N=150, while those with fewer than 100 subjects were excluded. This yielded a final cohort of 558 HC and 1,143 patients distributed across diagnostic subgroups in a relatively balanced manner: ADHD, ASD, ASD+ADHD, ADHD+ANX, ADHD+LD (each N = 150), ANX (N = 148), LD (N = 140), and ADHD+ANX+LD (N = 105). The HC group was further split into a training set (N = 447) and held-out test set (N = 111). Participant recruitment, inclusion criteria, and ethical approvals were handled locally by each contributing site, with informed consent obtained from all participants or their legal guardians. The reader can refer to Fig. 1 for a detailed breakdown of the final sample distribution across age, sex, site, diagnosis, and comorbidity. The original sample distribution before resampling is shown in Supplementary Fig. S1. A comprehensive overview of the aggregated datasets—including participant demographics, data collection sites, selection and diagnostic criteria, and available medication information—is provided in Supplementary Data 1.

### EEG Acquisition and Preprocessing

Across datasets, high-density (128-channel) resting-state EEG were recorded using the 128-channels EGI system (Electrical Geodesics, Inc.). For the present study, we exclusively analyzed data from the eyes-open condition. Across all sites, EEG acquisition protocols followed standardized procedures suitable for pediatric and clinical populations. EEG preprocessing and artifact removal pipeline was performed using a multi-stage, automated pipeline, supported by visual inspection. First, raw EEG signals were bandpass filtered between 1 and 100 Hz, and downsampled to a common sampling rate of 200 Hz. Bad EEG channels were identified using the ‘*pyprep*’ algorithm, which employs a *RANSAC*-based procedure to identify outlier channels by comparing them to model-predicted signals derived from randomly selected subsets of channels^55^. Identified bad channels were subsequently interpolated using data from neighboring electrodes. Next, re-referencing was performed using the common average reference method to minimize common noise across electrodes. Independent Component Analysis (ICA) was then applied and the ‘*IClabel*’ algorithm automatically identified components related to eye blinks^56^. A second bandpass filter was then applied to narrow the frequency range to [1-45Hz]. Next, EEG signals were segmented into 10-second epochs and the ‘*Autoreject*’ toolbox^57^ was used to detect and correct/reject artifactual epochs. All datasets underwent the preprocessing steps described above, unless a specific step was not feasible for a given dataset. For example, the Autoreject step was omitted for the ABCCT dataset, and bad channel detection and interpolation were not applied to the femaleASD dataset. Subjects were excluded if they met any of the following criteria: (i) more than 20% of channels required interpolation, or (ii) they were flagged during final visual inspection as having poor data quality. Finally, EEG signals were normalized within each subject by subtracting the mean and dividing by the standard deviation.

### Multidimensional EEG Features

We first reduced the spatial resolution of the preprocessed data by downsampling from 128 to 19 standard EEG channels, aligned with the widely adopted ‘*Nihon Kohden EEG 2100*’ system.

In total, we extracted 1,957 EEG features per subject (103 features × 19 channels), spanning four key categories: Catch24, Complexity, Spectral, and Time-Domain Features. Catch24 features are derived from the Canonical Time-series CHaracteristics set and summarize signal dynamics using statistical and autocorrelation properties^58^. Complexity features capture measures of entropy, fractal dimension, and non-linear dynamics across frequency bands. Spectral features quantify oscillatory power and aperiodic (1/f-like) components of the signal. Time-domain features characterize waveform shape and amplitude characteristics, including skewness, kurtosis, and total power, among others. Together, these features provide a comprehensive, multi-dimensional representation of brain activity. See Fig. 2 for an illustration of the number and distribution of features across categories. A complete overview of all EEG features used in this study is provided in Supplementary Data 2.

To account for potential site-related variability in the extracted EEG features, we applied the NeuroCombat harmonization^22^ (https://github.com/Jfortin1/neuroCombat), using age, sex and diagnostic group as covariates.

### Normative Modeling

To quantify individual deviations in EEG features relative to a normative population, we implemented normative modeling using the Generalized Additive Models for Location, Scale, and Shape (GAMLSS) framework (‘*gamlss R*’ package)^59^. This approach models each EEG feature as a function of age, while accommodating nonlinearity and heteroscedasticity.

For each feature, we first identified the best-fitting distribution family and covariate structure. An empirical model selection procedure was employed: GAMLSS models were trained across a broad set of continuous and mixed distribution families (with ≥3 moments), and sex was included as an additional covariate. The model with the lowest Bayesian Information Criterion (BIC) was selected and used to derive normative curves. Then, model performance was evaluated via convergence diagnostics, normalized quantile residuals, and normal Q-Q plots.

GAMLSS models were trained on the healthy control training set (HC_tr; 80% of healthy participants). These models were then applied to the clinical groups and the held-out healthy controls (HC_te; remaining 20%) to compute individualized deviation scores (z-scores). Deviation scores were derived from quantile randomized residuals^60^, providing a standardized metric of subject-specific deviations from the normative trajectory. This resulted in individualized deviation maps for each participant across all EEG features and channels.

### Transdiagnostic Clustering

To identify neurophysiologically distinct subgroups across the diagnostic spectrum, we implemented a data-driven unsupervised clustering approach using Similarity Network Fusion (SNF)^19^, a powerful integrative method for combining multimodal data into a unified similarity network. SNF constructs subject-by-subject similarity networks from each data source and iteratively fuses them into a consensus network, capturing both shared and complementary information across modalities.

In our study, we applied SNF to cluster participants based on their neurophysiological deviation profiles derived from normative modeling. Specifically, we divided deviation z-scores across all 19 EEG channels into four major feature domains—Catch24, complexity, spectral, and time-domain features—computed for all clinical groups and held-out healthy controls (total N = 1254). Each feature domain was treated as an independent data source for which the subjects similarity matrix was computed using squared Euclidean distance (‘*sqeucl*’), representing how similar individuals were to each other based on their deviation scores.

These domain-specific similarity matrices were then fused using the ‘*SNFpy*’ Python package (https://github.com/rmarkello/snfpy). Default SNF parameters were used (number of neighbors *K* = 20, hyperparameter *alpha* = 0.5, and number of iterations *T* = 20), which are known to offer stable performance across studies^19^.

Next, spectral clustering was applied to the fused similarity matrix. Spectral clustering partitions the network by minimizing edge weights between clusters, effectively grouping individuals with high intra-cluster similarity. To determine the optimal number of clusters, we systematically tested solutions ranging from 2 to 20 clusters, evaluating each using two established criteria: the ‘*eigen-gap*’ method^61^ and ‘*rotation cost*’ minimization^62^. These criteria identified an optimal solution that effectively separated individuals into clinically meaningful subgroups with distinct neurophysiological patterns.

Then, to evaluate the contribution of each data source and individual feature to the final clustering solution, we calculated ‘*Normalized Mutual Information (NMI)*’ scores, which measure the agreement between two clustering assignments, ranging from 0 (no agreement) to 1 (perfect agreement)^27,63^. Specifically, we calculated NMI between the clustering derived from each individual feature domain and the final fused clustering outcome from SNF, thereby assessing their relative contribution to the integrated clustering structure^64^. We also computed feature-specific NMI scores, measuring the similarity between the clustering based on each individual feature and the overall SNF-derived solution. Higher NMI scores reflect greater influence or concordance of the domain or feature with the final cluster assignment. These analyses offer insights into which features and data modalities most strongly drive the transdiagnostic clustering structure.

Finally, to evaluate the robustness of the identified clusters under small perturbations of the data, we performed a subsampling-based stability analysis^65^. Specifically, we repeated the entire SNF clustering pipeline 100 times, each time randomly removing 10% of the participants (i.e., 125 subjects). At each iteration, we computed a robustness coefficient by comparing the obtained cluster assignments of the retained subjects to their assignments in the main full-sample clustering, using the adjusted ‘*Rand index*’ (ARI)^66^. As an additional validation step, we assessed potential confounding effects of key covariates on cluster membership. For continuous variables (age and comorbidity), we used the Mann–Whitney U test with Bonferroni correction (p < 0.01), and quantified effect sizes using Cohen’s d (d > 0.5 indicating moderate effects). For categorical variables (sex and acquisition site), we applied Chi-squared tests with Bonferroni correction (p < 0.01), and reported effect sizes using Cramér’s V (V > 0.3 indicating moderate effects).

### Behavioral Measures

We selected common behavioral scores across datasets and grouped them into 13 functionally and clinically meaningful behavioral domains, including Hyperactivity, Attention, Depression, Aggression, Internalization, Externalization, Anxiety, Social Functioning, Autistic Traits, Family Conflict, Executive Function, Cognitive Function, Language, and Emotional Reactivity. This domain-based approach reduced redundancy from multiple instruments assessing similar constructs and facilitated more consistent interpretation of behavioral profiles across sites. To derive a single behavioral indicator per domain, we computed composite z-scores. Specifically, raw scores within each domain were first standardized across the entire cohort (mean-centered and scaled by the standard deviation), and then averaged to yield a domain-level composite^67–69^. The full list of behavioral domains and their constituent scores is provided in Supplementary Table S2.

### Clinical Analysis

Following the identification of neurophysiological clusters, we conducted statistical analyses to examine whether behavioral domains differed significantly across clusters. We first applied the non-parametric Kruskal–Wallis H test^70^ to detect behavioral measures that significantly discriminated between clusters. Then, for behaviors showing significant overall effects (p < 0.05), we conducted pairwise post hoc comparisons using Dunn’s test with Bonferroni correction^71^ to identify specific cluster differences. Finally, we performed Multiple Linear Regression (MLR) analyses using the built-in function in scikit-learn python package to further investigate the relationship between EEG features and clinical outcomes^72^. MLR linked feature coefficients to the most prominent behavioral traits within each cluster. These prominent behavioral traits were identified based on the mean composite z-score, with higher absolute values indicating more pronounced problem expression of that behavior in the corresponding cluster.

## Data Availability

ABCCT and femaleASD datasets can be requested from the NIMH Data Archive platform (https://nda.nih.gov/edit_collection.html?id=2288, https://nda.nih.gov/edit_collection.html?id=2021); MIPDB and HBN datasets are accessible at Child Mind Institute website (https://fcon_1000.projects.nitrc.org/indi/cmi_eeg/index.html, https://fcon_1000.projects.nitrc.org/indi/cmi_healthy_brain_network/index.html); LausanneASD dataset can be made available upon request.

## Code Availability

Codes are available at https://github.com/MINDIG-1/NM-Clinical. We used ‘*gamlss*’ R package^59^ for statistical modeling, ‘*MNE*’ python package (https://mne.tools/stable/index.html) for EEG signal processing, ‘*SNFpy*’ python package (https://github.com/rmarkello/snfpy) for SNF clustering, and custom python-based scripts for the remaining analysis and visualization.

## Acknowledgment

This work was funded by MINDIG as a part of its R&D activity. This work has been supported by a French government grant managed by the Agence Nationale de la Recherche under the France 2030 program, EEG-MIND (ANR-22-EXPR0009). It was also supported by the “Region Bretagne”, Inno R&D project no. 23001155 and Rennes Metropole (AICE project) and the INCR (PsyNorm and Creapark projects). We would like to thank all the researchers who shared their data in open-access and all the participants (patients and controls) who approved the use of their data in research.

## Authors Contribution

J.T., M.H., S.A., and A.E. conceived the study and oversaw data analysis and interpretation. J.T. drafted the manuscript and generated the figures with critical revisions from all authors. A.M. managed data access and organization. A.E. and S.A. assisted with data preprocessing and feature extraction. G.R., A.L., and A.I. contributed to behavioral analysis and interpretation. B.R. and N.C. provided access to ASD datasets.

## Ethics Declaration

### Ethics approval

Ethical oversight was provided by the Chesapeake Institutional Review Board, Yale Institutional Board (Yale, SCRI) and the *Commission Cantonale d’Ethique de la Recherche sur l’être humain* (CER-VD, Switzerland). All procedures performed were in accordance with the ethical standards of the 1964 Helsinki Declaration and its later amendments or comparable ethical standards. All participants (or their legal guardians) provided written informed consent prior to inclusion in the study.

## Supplementary information

Supplementary Data 1 (excel)

Supplementary Data 2 (excel)

Supplementary Data 3 (excel)

Supplementary Information (doc)

