## Supplementary Materials for "Transdiagnostic electrophysiological subtypes reveal brain-behavior dimensions in youth psychiatry"

**Supplementary Data 1: Demographic details, EEG system specifications, selection criteria, ethical approvals, and medication information for the five datasets included in this study.**

Supplementary Data 1 (Excel File)

**Supplementary Data 2: Detailed specifications of the EEG features used in this study.**

Supplementary Data 2 (Excel File)

**Supplementary Data 3: Normative models results, including the optimal family and parameters obtained for each EEG feature.**

Supplementary Data 3 (Excel File)

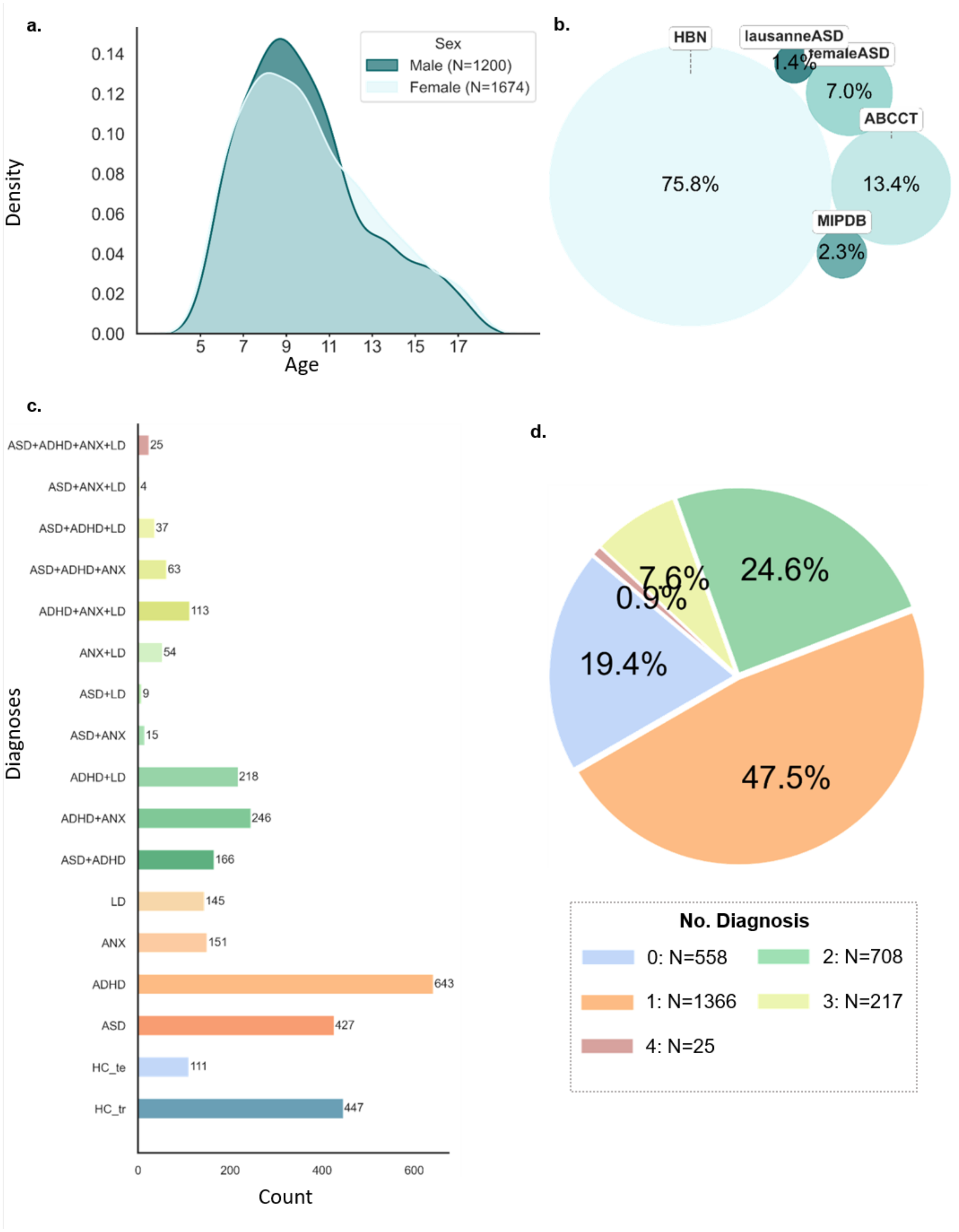


**Fig. S1: Overview of original data sample characteristics (before resampling) across age, sex, diagnostic groups, acquisition sites, and comorbidity profiles.**

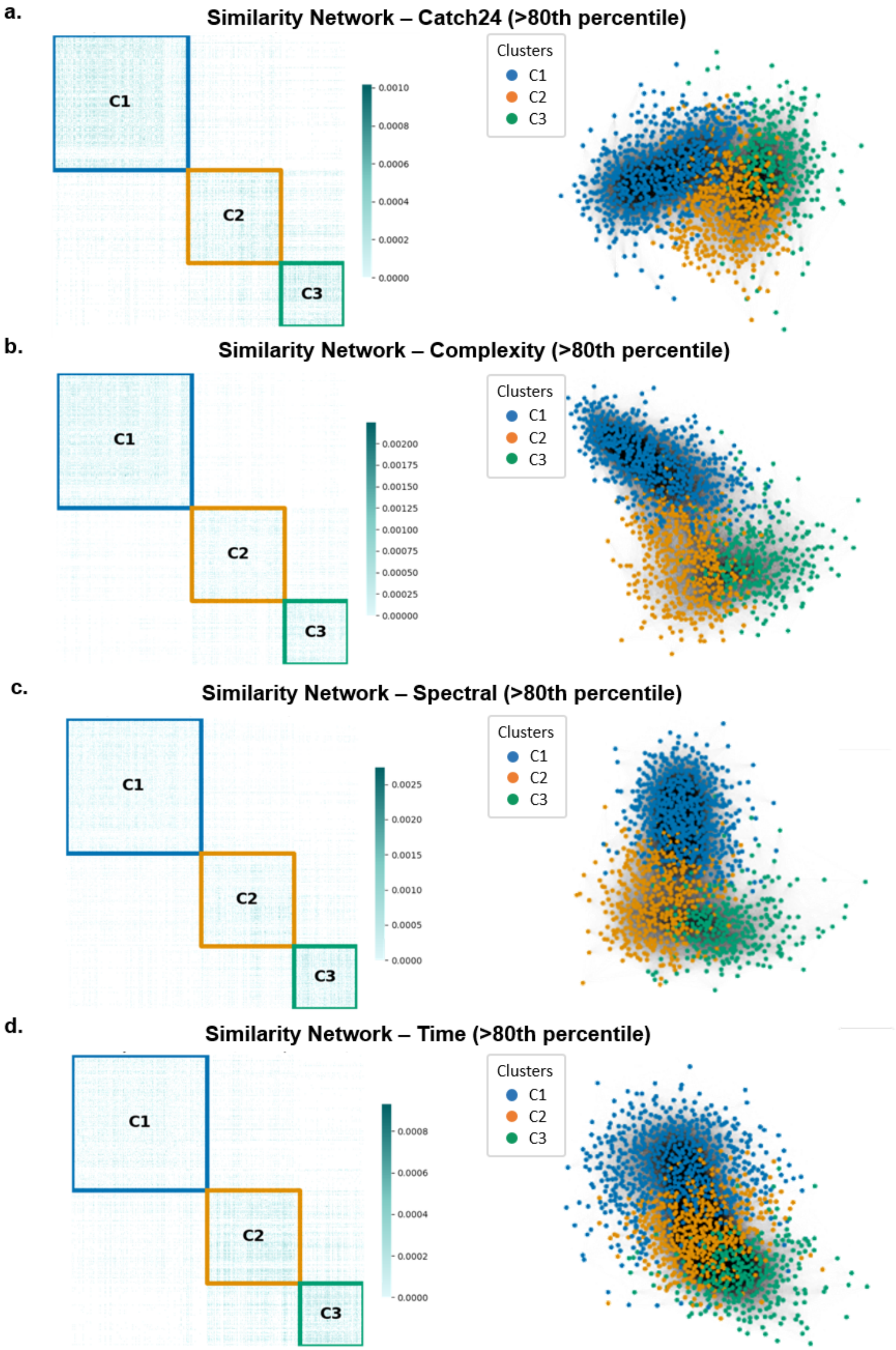
a.
Similarity Network – Catch24 (>80th percentile)
C1
C2
C3
0.0010
0.0008
0.0006
0.0004
0.0002
0.0000
Clusters
C1
C2
C3
b.
Similarity Network – Complexity (>80th percentile)
C1
C2
C3
0.00200
0.00175
0.00150
0.00125
0.00100
0.00075
0.00050
0.00025
0.00000
Clusters
C1
C2
C3
c.
Similarity Network – Spectral (>80th percentile)
C1
C2
C3
0.0025
0.0020
0.0015
0.0010
0.0005
0.0000
Clusters
C1
C2
C3
d.
Similarity Network – Time (>80th percentile)
C1
C2
C3
0.0008
0.0006
0.0004
0.0002
0.0000
Clusters
C1
C2
C3

**Fig. S2: Similarity networks per data source (feature category) visualized as heatmaps and graphs.** Illustrations are thresholded at the 80th percentile of the similarity strength. In the graph, nodes represent participants and edges represent pairwise similarity strength.

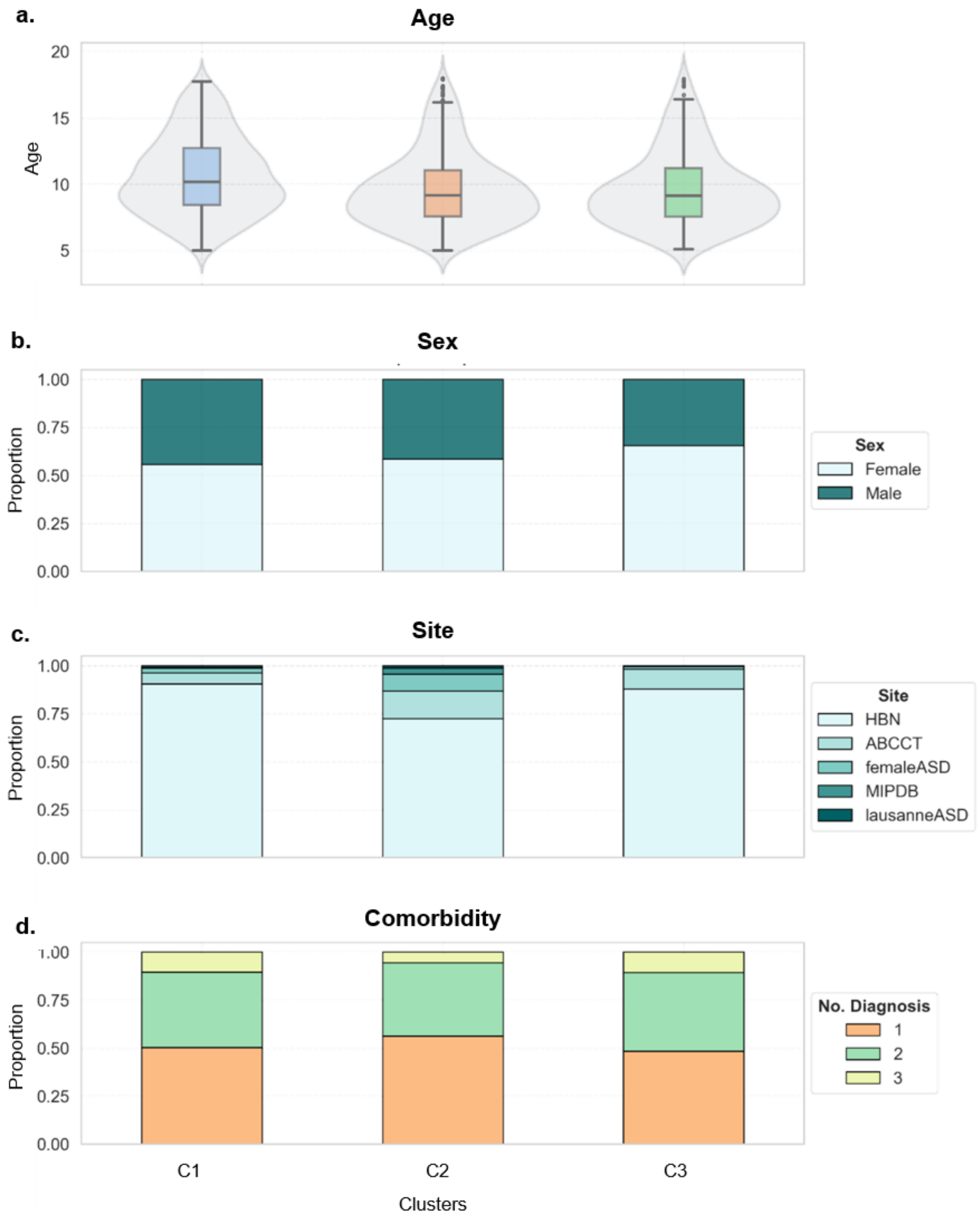


**Fig. S3: Covariate distributions across clusters.** Covariates include age, sex, acquisition sites, and comorbidity profiles.

**Table. S1: Statistical comparison of covariates (age, sex, site, and comorbidity) across clusters**

| | | Age | | Sex | | Site | | Comorbidity | |
|---|---|---|---|---|---|---|---|---|---|
| | | p_value (corr) | Effect Size | p_value (corr) | Effect Size | p_value (corr) | Effect Size | p_value (corr) | Effect Size |
| **C1** | **C2** | 1.02e-07 | 0.35 | 1.0 | 0.0 | 3.75e-12 | 0.03 | 0.09 | 0.17 |
| **C1** | **C3** | 3.00e-06 | 0.34 | 0.02 | 0.08 | 0.13 | 0.08 | 1.0 | -0.03 |
| **C2** | **C3** | 1.0 | -0.003 | 0.23 | 0.06 | 6.61-06 | 0.20 | 0.07 | -0.21 |

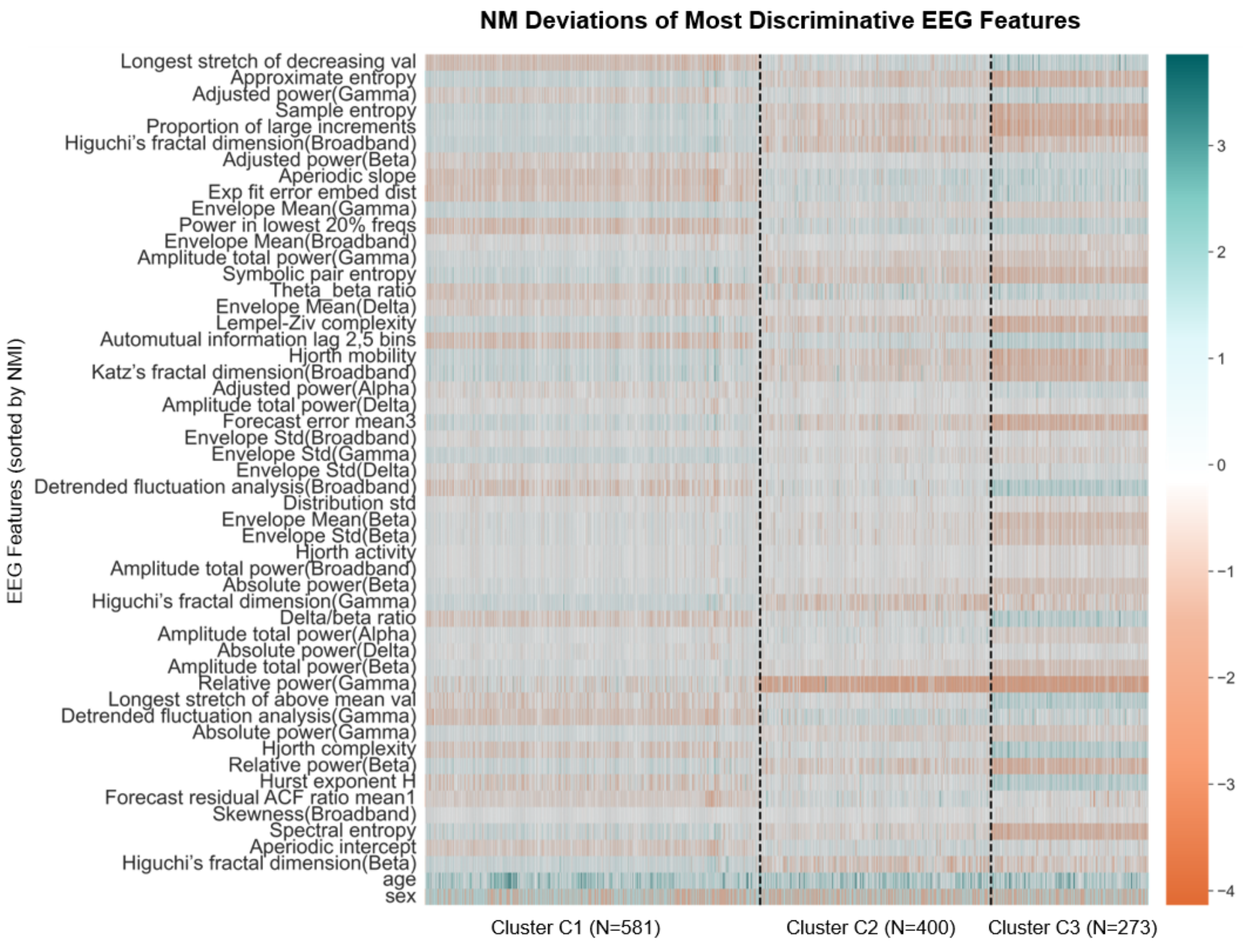


**Fig. S4: Z-score distribution of the top 50 EEG features (sorted by NMI scores; top to bottom) across clusters.** Green color indicates positive deviations and orange indicates negative deviations.

**Table. S2: Behavior scores categorized by domain**

| Behavior Domain | Behavior Scores |
|---|---|

| | |
|---|---|
| **Hyperactivity** | SWAN_Hyperactivity_Avg, C3SR_Hyperactivity_Raw, SDQ_Hyperactivity_Total |
| **Attention** | CBCL_Attention_Raw, SWAN_Inattention_Avg, C3SR_Inattention_Raw |
| **Depression** | CBCL_Withdrawn_Raw, MFQ_SR_Score, MFQ_P_Score, SDQ_Emotional_Problems_Total |
| **Aggression** | CBCL_Aggressive_Raw, C3SR_Aggression_Raw |
| **Internalization** | CBCL_Internal_Raw, SDQ_Internalising_Total |
| **Externalization** | CBCL_External_Raw, SDQ_Externalising_Total |
| **Anxiety** | CBCL_Anxious_Raw, SCARED_SR_Score, SCARED_P_Score, PANAS_NegativeAffect |
| **Social** | SRS_Communication_Raw, CBCL_Social_Raw, SDQ_Prosocial_Total |
| **AutisticTraits** | SCQ_Score, ASSQ_Score, SRS_Total_Raw |
| **FamilyConflict** | CPIC_Frequency_Total, CPIC_Intensity_Total, CPIC_Resolution_Total, CPIC_Content_Total, CPIC_Perceived_Threat_Total, CPIC_Self_Blame_Total, CPIC_Triangulation_Total, CPIC_Stability_Total, PSI_Score |
| **ExecutiveFunction** | NIH_final_Card_Sort_Raw, NIH_final_List_Sort_Raw, NIH_final_Processing_Raw |
| **CognitiveFunction** | WISC_SymbolSearch_Raw, WISC_Vocabulary_Raw, WISC_VCI_Raw, WISC_FullScale4_Raw, WISC_BlockDesign_Scaled, WISC_Similarities_Scaled, WISC_Matrix_Scaled |
| **Language** | CTOPP_Elision_Raw, CTOPP_BlendingWords_Raw, CTOPP_NonWordRepetition_Raw, CTOPP_RapidDigitNaming_Raw, CTOPP_RapidSymbolicNaming_Composite, TOWRE_Score_Raw |
| **Emotional** | NLES_P_Upset_Total |

**Table. S3: Kruskal-Wallis Test Results for Behavioral Domains Across Clusters. Significance levels are indicated as follows: ****p<0.05*, **p<0.01, ***p<0.001*.***

| Behavior | Cluster | Median | IQR | Mean | SD | H-stat | p-value |
|---|---|---|---|---|---|---|---|
| **Hyperactivity** | 1 | -0.0791 | 1.0562 | -0.0325 | 0.7772 | 18.8814 | 7.94e-05*** |
| | 2 | 0.0332 | 1.2788 | -0.0435 | 0.9062 | | |
| | 3 | 0.2480 | 1.1178 | 0.2179 | 0.7835 | | |
| **Attention** | 1 | 0.0538 | 1.1042 | 0.0314 | 0.7952 | 21.4238 | 2.23e-05*** |
| | 2 | -0.2045 | 1.3600 | -0.1977 | 0.8894 | | |
| | 3 | 0.1509 | 1.1250 | 0.1371 | 0.7588 | | |
| **Depression** | 1 | -0.0775 | 1.0318 | 0.0957 | 0.8241 | 21.8683 | 1.78e-05*** |
| | 2 | -0.3249 | 0.8689 | -0.1474 | 0.7177 | | |
| | 3 | -0.1859 | 0.8500 | -0.0152 | 0.7378 | | |
| **Aggression** | 1 | -0.2190 | 1.1870 | 0.0215 | 0.8587 | 4.7870 | 0.0913 |
| | 2 | -0.3036 | 1.1597 | -0.0536 | 0.9001 | | |
| | 3 | -0.1646 | 1.1521 | 0.0856 | 0.9220 | | |
| **Internalization** | 1 | -0.0757 | 1.3665 | 0.1178 | 0.9626 | 31.2402 | 1.65e-07*** |
| | 2 | -0.4467 | 1.1811 | -0.2200 | 0.8707 | | |
| | 3 | -0.0977 | 1.0700 | 0.0206 | 0.8484 | | |
| **Externalization** | 1 | -0.0986 | 1.2792 | -0.0004 | 0.9045 | 8.6660 | 0.0131** |
| | 2 | -0.2486 | 1.2607 | -0.0943 | 0.9332 | | |
| | 3 | 0.0473 | 1.1877 | 0.1012 | 0.9257 | | |
| **Anxiety** | 1 | -0.0708 | 1.0874 | 0.0801 | 0.7838 | 24.4594 | 4.88e-06*** |
| | 2 | -0.3065 | 0.9878 | -0.1693 | 0.7207 | | |
| | 3 | -0.0997 | 0.9019 | 0.0251 | 0.7355 | | |
| **Social** | 1 | -0.1167 | 1.2130 | 0.0562 | 0.8733 | 11.3014 | 0.0035*** |
| | 2 | -0.2726 | 1.2250 | -0.0551 | 0.9360 | | |
| | 3 | 0.0290 | 1.3421 | 0.1500 | 0.9252 | | |
| **Autistic Traits** | 1 | -0.1899 | 1.1029 | 0.0424 | 0.9249 | 9.3071 | 0.0095*** |
| | 2 | -0.3073 | 1.1873 | -0.0122 | 1.0173 | | |
| | 3 | -0.1124 | 1.1027 | 0.1368 | 0.9179 | | |
| **Family Conflict** | 1 | 0.0792 | 0.9185 | 0.0156 | 0.7849 | 0.0768 | 0.9622 |
| | 2 | 0.0796 | 0.9300 | 0.0321 | 0.8142 | | |
| | 3 | 0.0226 | 0.8500 | 0.0451 | 0.6768 | | |
| **Executive Function** | 1 | 0.1566 | 0.7853 | 0.1014 | 0.7201 | 23.1585 | 9.36e-06*** |
| | 2 | 0.0102 | 0.9436 | -0.1125 | 0.7726 | | |
| | 3 | 0.0092 | 0.8098 | -0.1183 | 0.6917 | | |

| | | | | | | | |
|---|---|---|---|---|---|---|---|
| **Cognitive Function** | 1 | 0.0161 | 1.0434 | -0.0004 | 0.7361 | 0.0244 | 0.9878 |
| | 2 | -0.0695 | 1.0536 | 0.0044 | 0.7751 | | |
| | 3 | -0.0170 | 1.0377 | -0.0087 | 0.7395 | | |
| **Language** | 1 | 0.0225 | 0.6905 | 0.0161 | 0.5236 | 1.6557 | 0.4369 |
| | 2 | -0.0434 | 0.6623 | -0.0230 | 0.4984 | | |
| | 3 | 0.0086 | 0.6624 | 0.0127 | 0.5246 | | |
| **Emotional** | 1 | -0.0713 | 1.3725 | 0.1420 | 1.0848 | 16.7804 | 0.0002*** |
| | 2 | -0.2543 | 1.0980 | -0.1619 | 0.8793 | | |
| | 3 | -0.3458 | 1.2123 | -0.1049 | 0.9090 | | |

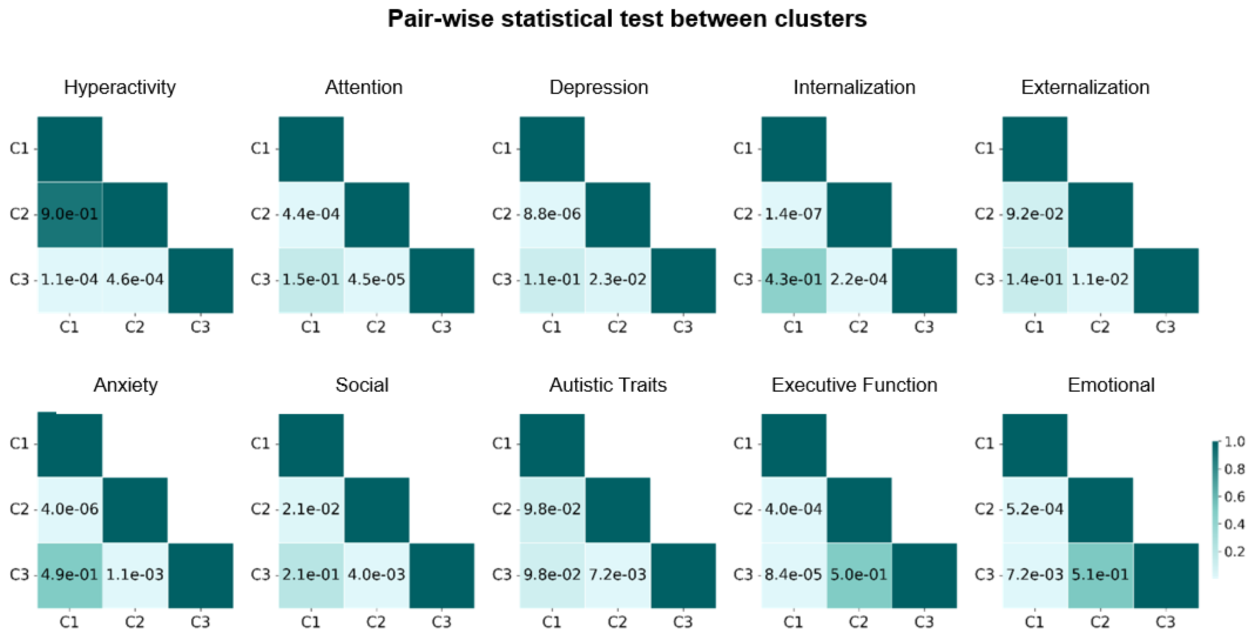


**Fig. S5: P-values Results of Dunn's pairwise test applied on each behavioral domain to identify inter-cluster differences.**

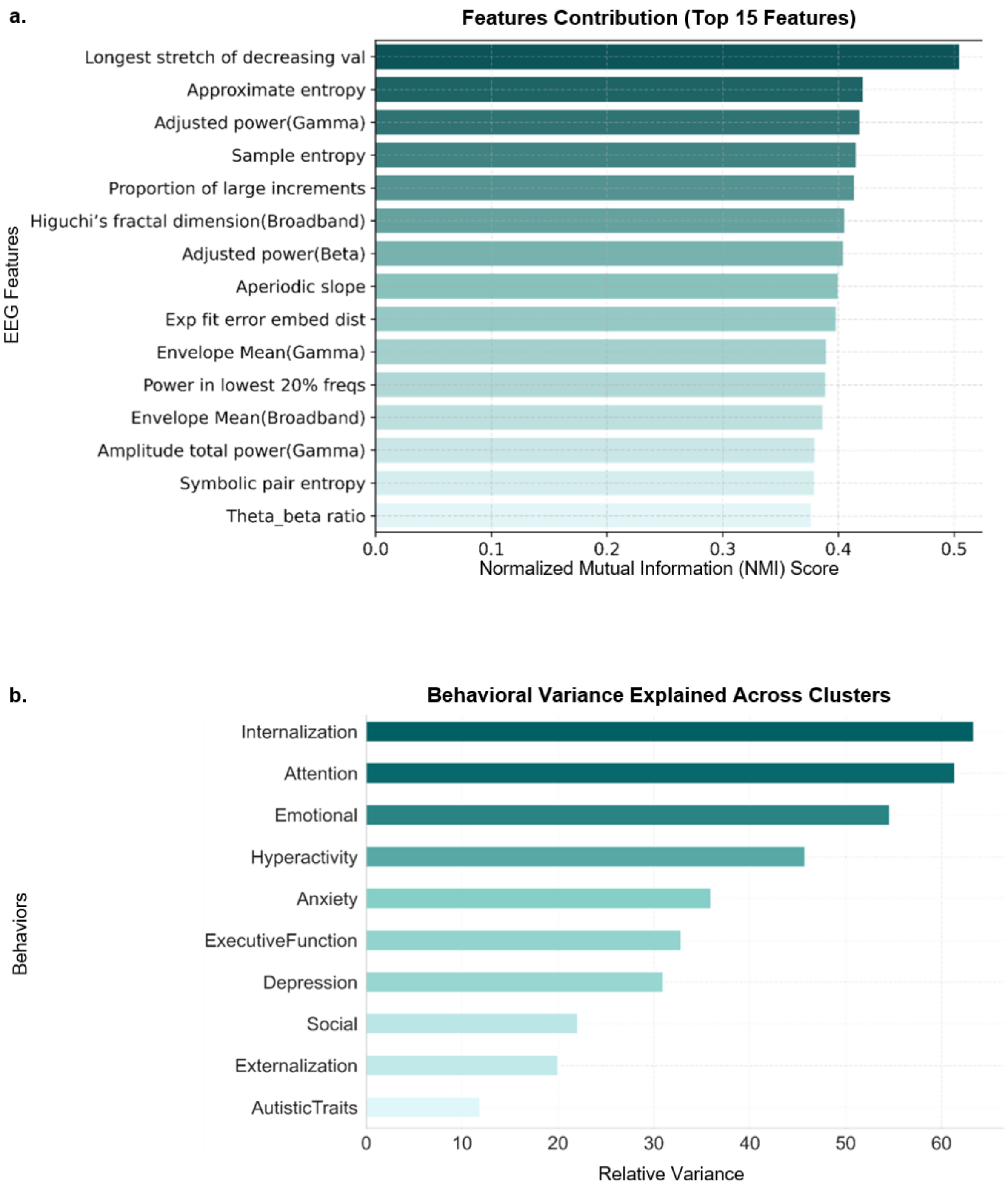


**Fig. S6: Electrophysiological and behavioral drivers of clustering. a** Feature contributions to SNF clustering of the top 15 features (sorted by NMI scores). **b** Behavioral domain contributions to the differentiation between clusters (sorted by explained variance values).

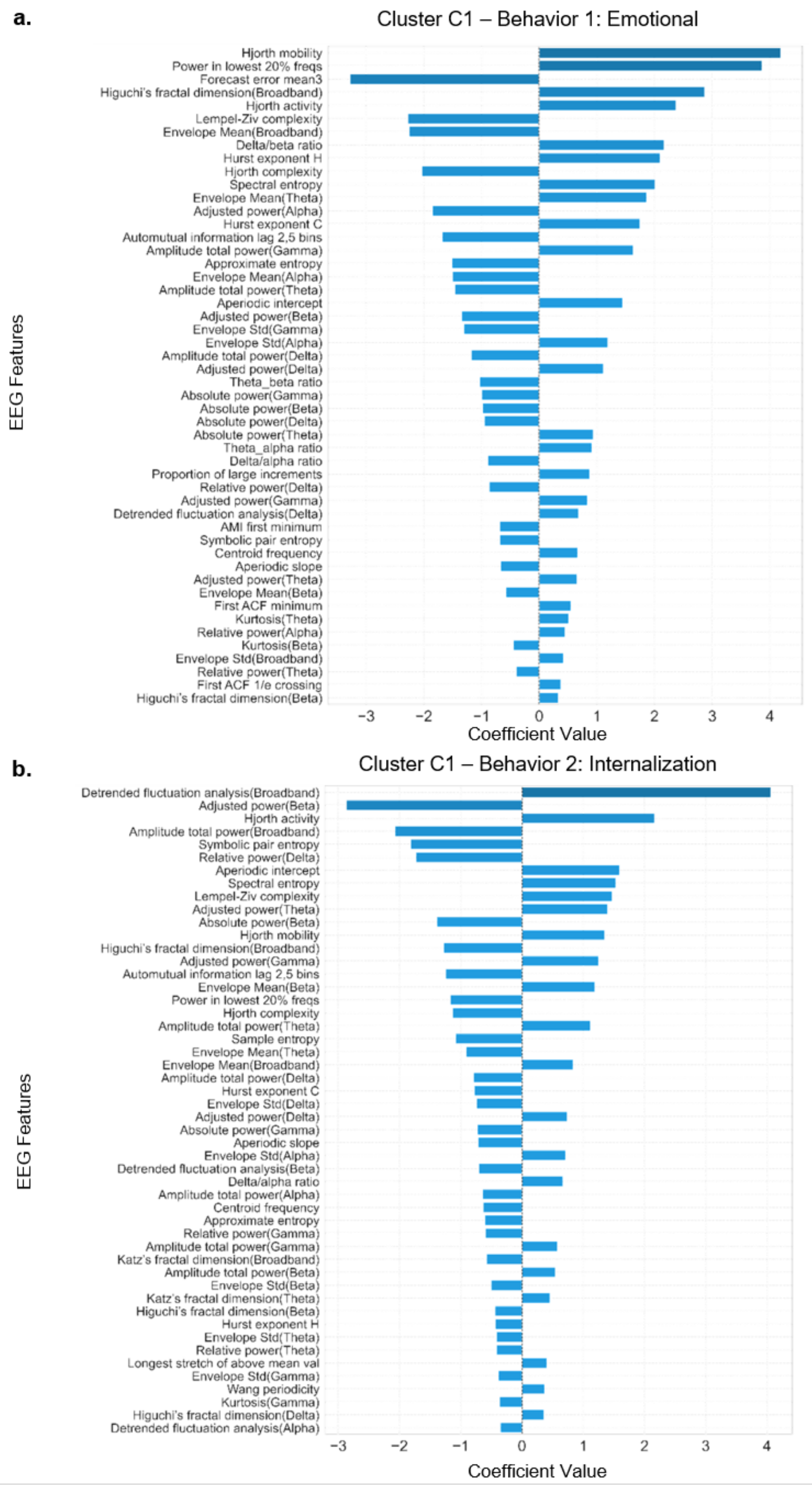


**Fig. S7: Standardized regression coefficients for the top 50 EEG features contributing to the prediction of the two most prominent behavioral traits of cluster C1.** Features are ranked from highest (top) to lowest (bottom) absolute values.

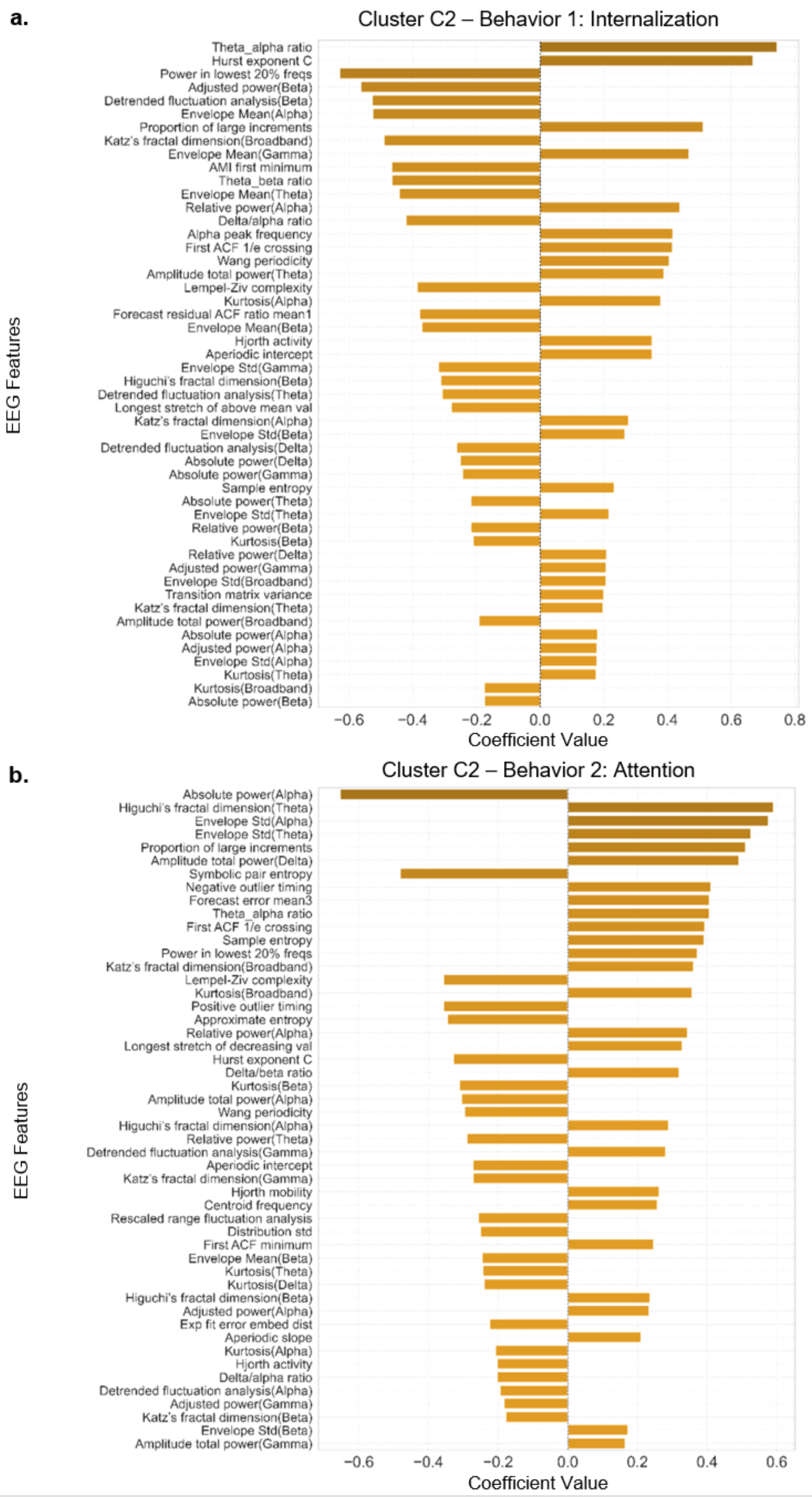


**Fig. S8: Standardized regression coefficients for the top 50 EEG features contributing to the prediction of the two most prominent behavioral traits of cluster C2.** Features are ranked from highest (top) to lowest (bottom) absolute values.

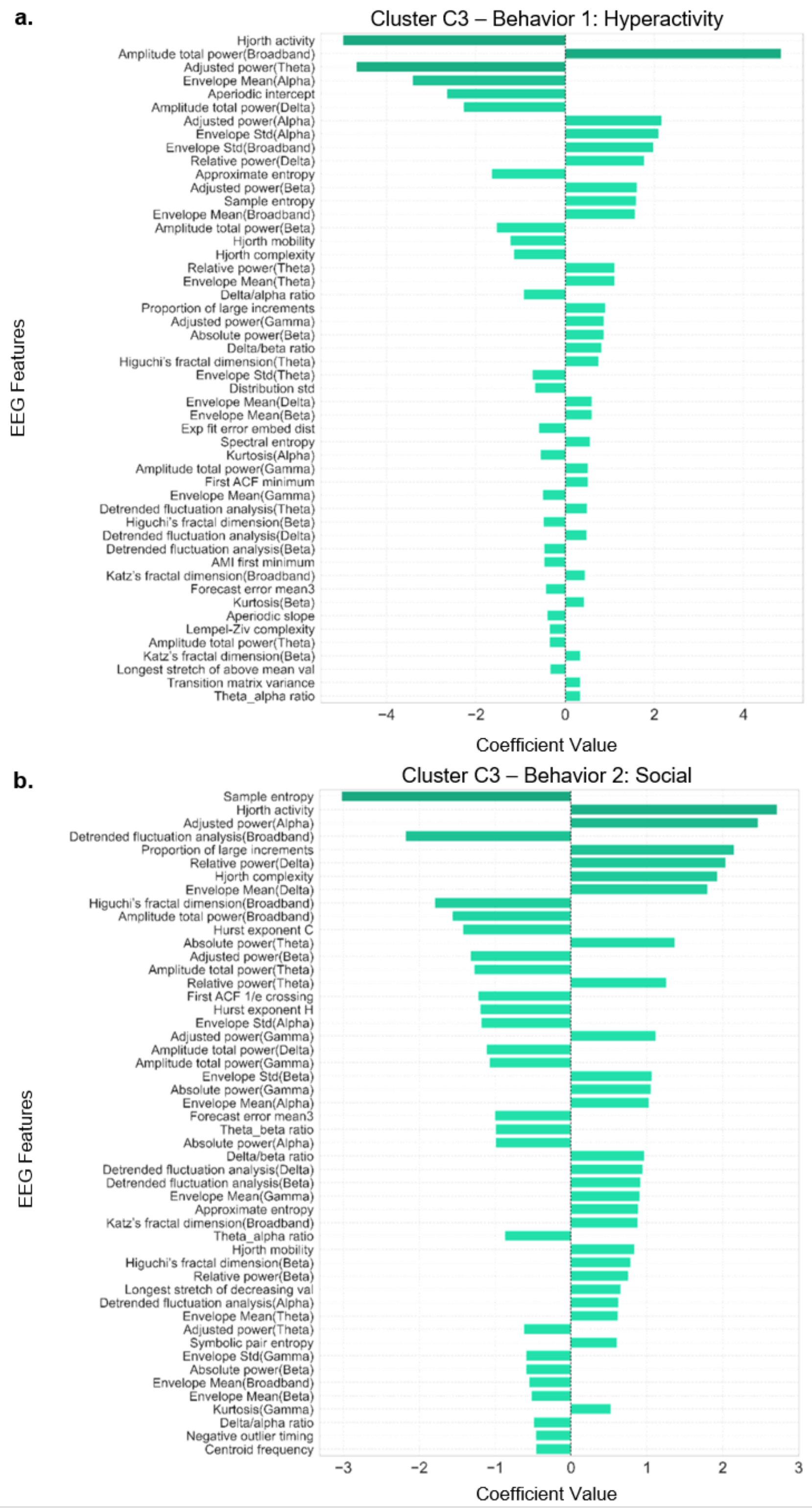


**Fig. S9: Standardized regression coefficients for the top 50 EEG features contributing to the prediction of the two most prominent behavioral traits of Cluster c3.** Features are ranked from highest (top) to lowest (bottom) absolute values.
