## Supplementary Data 1 for "Transdiagnostic electrophysiological subtypes reveal brain-behavior dimensions in youth psychiatry"

| Dataset | Groups | HC | | Patients (original, before resampling) | | Patients (final, after resampling) | | EEG Recording System | | | Site | Source | Download Reference | Diagnostic/Selection Criteria | Medication Information |
|---|---|---|---|---|---|---|---|---|---|---|---|---|---|---|---|
| | | No subjs (% Male) | Age (year): median (SD) [range] | No subjs (% Male) | Age (year): median (SD) [range] | No subjs (% Male) | Age (year): median (SD) [range] | Montage | Number of channels | Sampling frequency | | | | | |
| **ABCCT (Autism Biomarkers Consortium for Clinical Trials)** | Healthy Controls (HC) | 118 (69.5%) | 8.4 (1.6) [6.1–11.5] | | | | | EGI128 | 128 downsampled to 19 | 1000 | , Boston Children's Hospital (BCH), University of Washington/Seattle Children's Re | Open (NIMH Data Archive) | ://nda.nih.gov/edit_collection.html?id= | n the Autism Diagnostic Observation Schedule (ADOS) and the A | ired for 8 weeks prior to enrollment: All medications were allowed in order to enroll a representative samp |
| | Autism Spectrum Disorder (ASD) | | | 268 (76.9) | 8.3 (1.6) [6.1–11.6] | 94 (77.7) | 8.5 (1.6) [6.1–11.5] | | | | | | | | |
| **femaleASD (Multimodal Developmental Neurogenetics of Females with ASD)** | Healthy Controls (HC) | 101 (53.5) | 13.2 (2.8) [8.1–17.9] | | | | | EGI128 | 128 downsampled to 19 | 500 | Children's Hospital (BCH), University of Washington/Seattle Children's Research I | Open (NIMH Data Archive) | ://nda.nih.gov/edit_collection.html?id= | h additional parent report using the Social Responsiveness Scale, | epine, barbiturate, or anti-epileptic medication, or participants with medication changes within the 6 weeks |
| | Autism Spectrum Disorder (ASD) | | | 101 (52.5) | 12.2 (2.9) [8.0–17.8] | 35 (48.6) | 11.8 (2.8) [8.2–17.3] | | | | | | | | |
| **LausanneASD** | Healthy Controls (HC) | 29 (37.9) | 6.7 (1.0) [5.2–9.0] | | | | | EGI128 | 128 downsampled to 19 | 250/1000 | Lausanne, SwitzerlandLausanne University Hospital (STSA, CHUV) | Not Open | - | *Diagnosis*: DSM-5 diagnosis of ASD based on ADOS and ADI-R | - |
| | Autism Spectrum Disorder (ASD) | | | 12 (83.3) | 6.3 (1.4) [5.0–9.1] | 6 (100.0) | 6.6 (1.7) [5.0–9.1] | | | | | | | | |
| **MIPDB (Multimodal Resource for Studying Information Processing in the Developing Brain)** | Healthy Controls (HC) | 59 (61.0) | 13.0 (3.0) [8.0–18.0] | | | | | EGI128 | 128 downsampled to 19 | 500 | New York CityChild Mind Medical Practice | Open (Child Mind Institute) | _1000.projects.nitrc.org/indi/cmi_eeg | *tion*: No contraindications for EEG (e.g., history of seizures or epil | - |
| | Attention deficit hyperactivity disorder (ADHD) | | | 8 (75.0) | 11.0 (2.3) [8.0–14.0] | 2 (50.0) | 10.5 (3.5) [8.0–13.0] | | | | | | | | |
| **HBN (Healthy Brain Network)** | Healthy Controls (HC) | 251 (46.2) | 9.1 (3.0) [5.0–17.3] | | | | | EGI128 | 128 downsampled to 19 | 500 | New York City(Diverse Communities) | Open (Child Mind Institute, INDI) | jects.nitrc.org/indi/cmi_healthy_brain | ADHD System, Strengths and Weaknesses of ADHD Symptoms | n cognitive and behavioral testing, as well as functional brain mapping. Participants who choose not to disc |
| | Autism Spectrum Disorder (ASD) | | | 46 (26.1) | 8.7 (3.7) [5.3–17.5] | 15 (13.3) | 7.3 (3.9) [5.3–16.0] | | | | | | | | |
| | Attention deficit hyperactivity disorder (ADHD) | | | 635 (26.8) | 9.1 (2.9) [5.0–17.8] | 148 (29.7) | 9.1 (3.1) [5.1–17.8] | | | | | | | | |
| | Anxiety (ANX) | | | 151 (57.0) | 10.3 (3.1) [5.4–17.6] | 148 (56.1) | 10.3 (3.1) [5.4–17.6] | | | | | | | | |
| | Learning Disorder (LD) | | | 145 (48.3) | 9.5 (2.7) [5.0–17.3] | 140 (47.1) | 9.5 (2.7) [5.0–17.3] | | | | | | | | |
| | ASD+ADHD | | | 166 (12.0) | 8.9 (3.2) [5.1–17.9] | 150 (11.3) | 9.1 (3.2) [5.1–17.9] | | | | | | | | |
| | ADHD+ANX | | | 246 (39.8) | 10.2 (2.8) [5.5–17.7] | 150 (38.7) | 10.4 (2.7) [5.5–17.7] | | | | | | | | |
| | ADHD+LD | | | 218 (30.3) | 9.2 (2.7) [5.7–17.9] | 150 (29.3) | 9.2 (2.8) [5.7–17.9] | | | | | | | | |
| | ADHD+ANX+LD | | | 113 (43.4) | 10.1 (3.0) [5.8–17.7] | 105 (41.0) | 10.1 (2.9) [5.8–17.7] | | | | | | | | |
| | ASD+ANX | | | 15 (33.3) | 11.8 (4.1) [5.2–17.9] | | | | | | | | | | |
| | ASD+LD | | | 9 (11.1) | 9.4 (3.0) [7.7–17.0] | | | | | | | | | | |
| | ANX+LD | | | 54 (46.3) | 9.2 (2.9) [6.5–17.2] | | | | | | | | | | |
| | ASD+ADHD+ANX | | | 63 (19.0) | 10.4 (3.1) [5.5–17.6] | | | | | | | | | | |
| | ASD+ADHD+LD | | | 37 (18.9) | 10.0 (2.6) [6.1–15.6] | | | | | | | | | | |
| | ASD+ANX+LD | | | 4 (0.0) | 14.5 (4.1) [8.8–17.8] | | | | | | | | | | |
| | ASD+ADHD+ANX+LD | | | 25 (20.0) | 11.3 (2.8) [6.7–16.8] | | | | | | | | | | |
