## Supplementary Data 2 for "Transdiagnostic electrophysiological subtypes reveal brain-behavior dimensions in youth psychiatry"

| | Feature Sub-Category | Feature Name | Variable Name | Bands | Channels |
|---|---|---|---|---|---|
| **Catch24 (CAnonical Time-series CHaract** | Distribution shape | 5-bin histogram mode | DN_HistogramMode_5 | - | 19 |
| | Distribution shape | 10-bin histogram mode | DN_HistogramMode_10 | - | 19 |
| | Extreme event timing | Positive outlier timing | DN_OutlierInclude_p_001_mdrmd | - | 19 |
| | Extreme event timing | Negative outlier timing | DN_OutlierInclude_n_001_mdrmd | - | 19 |
| | Linear autocorrelation | First ACF 1/e crossing | CO_f1ecac | - | 19 |
| | Linear autocorrelation | First ACF minimum | CO_FirstMin_ac | - | 19 |
| | Linear autocorrelation | Power in lowest 20% freqs | SP_Summaries_welch_rect_area_5_1 | - | 19 |
| | Linear autocorrelation | Centroid frequency | SP_Summaries_welch_rect_centroid | - | 19 |
| | Simple forecasting | Forecast error mean3 | FC_LocalSimple_mean3_stderr | - | 19 |
| | Incremental differences | Forecast residual ACF ratio mean1 | FC_LocalSimple_mean1_tauresrat | - | 19 |
| | Incremental differences | Proportion of large increments | MD_hrv_classic_pnn40 | - | 19 |
| | Symbolic | Longest stretch of above mean val | SB_BinaryStats_mean_longstretch1 | - | 19 |
| | Symbolic | Longest stretch of decreasing val | SB_BinaryStats_diff_longstretch0 | - | 19 |
| | Symbolic | Symbolic pair entropy | SB_MotifThree_quantile_hh | - | 19 |
| | Non-linear autocorrelation | Automutual information lag 2,5 bins | CO_HistogramAMI_even_2_5 | - | 19 |
| | Non-linear autocorrelation | Time reversibility | CO_trev_1_num | - | 19 |
| | Linear autocorrelation | AMI first minimum | IN_AutoMutualInfoStats_40_gaussian_fmmi | - | 19 |
| | Symbolic | Transition matrix variance | SB_TransitionMatrix_3ac_sumdiagcov | - | 19 |
| | Linear autocorrelation | Wang periodicity | PD_PeriodicityWang_th0_01 | - | 19 |
| | - | Exp fit error embed dist | CO_Embed2_Dist_tau_d_expfit_meandiff | - | 19 |
| | Low-scale scaling | Rescaled range fluctuation analysis | SC_FluctAnal_2_rsrangefit_50_1_logi_prop_r1 | - | 19 |
| | Low-scale scaling | Detrended fluctuation analysis | SC_FluctAnal_2_dfa_50_1_2_logi_prop_r1 | - | 19 |
| | - | Distribution mean | DN_Mean | - | 19 |
| | - | Distribution std | DN_Spread_Std | - | 19 |
| **Complexity** | Entropy parameter | Approximate entropy | app_ent | - | 19 |
| | - | Detrended fluctuation analysis | detrend_fluc | 6 (broadband, delta, theta, alpha, beta, gamma) | 19 |
| | - | Higuchi's fractal dimension | higuchi_frac | 6 (broadband, delta, theta, alpha, beta, gamma) | 19 |
| | Hjorth parameter | Hjorth activity | hjorth_activity | - | 19 |
| | Hjorth parameter | Hjorth complexity | hjorth_complexity | - | 19 |
| | Hjorth parameter | Hjorth mobility | hjorth_mobility | - | 19 |
| | Hurst exponent parameter | Hurst exponent C | hurst_exp_c | - | 19 |
| | Hurst exponent parameter | Hurst exponent H | hurst_exp_h | - | 19 |
| | - | Katz's fractal dimension | katz_frac | 6 (broadband, delta, theta, alpha, beta, gamma) | 19 |
| | - | Lempel-Ziv complexity | lziv | - | 19 |
| | Entropy parameter | Sample entropy | samp_ent | - | 19 |
| | Entropy parameter | Spectral entropy | spect_ent | - | 19 |
| **Spectral** | Power parameter | Adjusted power | adjusted_power | 5 (delta, theta, alpha, beta, gamma) | 19 |
| | - | Aperiodic intercept | aperiodic_intercept | - | 19 |
| | - | Aperiodic slope | aperiodic_slope | - | 19 |
| | Power Bands ratio | Delta/alpha ratio | delta_alpha_ratio | - | 19 |
| | Power Bands ratio | Delta/beta ratio | delta_beta_ratio | - | 19 |
| | - | Alpha peak frequency | peak_alpha_freq | - | 19 |
| | Power parameter | Absolute power | power_abs | 5 (delta, theta, alpha, beta, gamma) | 19 |
| | Power parameter | Relative power | power_rel | 5 (delta, theta, alpha, beta, gamma) | 19 |
| | Power Bands ratio | Theta_alpha ratio | theta_alpha_ratio | - | 19 |
| | Power Bands ratio | Theta_beta ratio | theta_beta_ratio | - | 19 |
| **Time** | - | Envelope Mean | env_mean | 6 (broadband, delta, theta, alpha, beta, gamma) | 19 |
| | - | Envelope Std | env_std | 6 (broadband, delta, theta, alpha, beta, gamma) | 19 |
| | - | Kurtosis | kurt_t | 6 (broadband, delta, theta, alpha, beta, gamma) | 19 |
| | - | Amplitude total power | power_t | 6 (broadband, delta, theta, alpha, beta, gamma) | 19 |
| | - | Skewness | skew_t | 6 (broadband, delta, theta, alpha, beta, gamma) | 19 |

broadband': [1, 45]'delta': [1, 4]'theta': [4, 8]'alpha': [8, 13]'beta': [13, 30]'gamma': [30, 45]
