## Supplementary Data 3 for "Transdiagnostic electrophysiological subtypes reveal brain-behavior dimensions in youth psychiatry"

| category | Feature Variable Name | Distribution Family | mu_formula | si_formula | nu_formula | tau_formula |
|---|---|---|---|---|---|---|
| **catch24** | CO_Embed2_Dist_tau_d_expfit_meandiff | exGAUS | y ~ 1 + fp(age, 1) | ~1 | ~1 | ~1 |
| | CO_f1ecac | exGAUS | y ~ 1 + fp(age, 1) | ~1 | ~1 | ~1 |
| | CO_FirstMin_ac | ST3 | y ~ 1 + fp(age, 1) | ~1 | ~1 | ~1 |
| | CO_HistogramAMI_even_2_5 | -SEP3 | y ~ 1 + fp(age, 1) | ~1 | ~1 | ~1 |
| | CO_trev_1_num | ST4 | y ~ 1 + fp(age, 1) | ~1 | ~1 | ~1 |
| | DN_HistogramMode_10 | PE | y ~ 1 + fp(age, 1) | ~1 | ~1 | ~1 |
| | DN_HistogramMode_5 | PE | y ~ 1 + fp(age, 1) | ~1 | ~1 | ~1 |
| | DN_Mean | PE | y ~ 1 + fp(age, 1) | ~1 | ~1 | ~1 |
| | DN_OutlierInclude_n_001_mdrmd | PE | y ~ 1 + fp(age, 1) | ~1 | ~1 | ~1 |
| | DN_OutlierInclude_p_001_mdrmd | PE | y ~ 1 + fp(age, 1) | ~1 | ~1 | ~1 |
| | DN_Spread_Std | ST3 | y ~ 1 + fp(age, 1) | ~1 | ~1 | ~1 |
| | FC_LocalSimple_mean1_tauresrat | SHASHo | y ~ 1 + fp(age, 1) | ~1 | ~1 | ~1 |
| | FC_LocalSimple_mean3_stderr | GG | y ~ 1 + fp(age, 1) | ~1 | ~1 | ~1 |
| | IN_AutoMutualInfoStats_40_gaussian_fmmi | PE | y ~ 1 + fp(age, 1) | ~1 | ~1 | ~1 |
| | MD_hrv_classic_pnn40 | ST3 | y ~ 1 + fp(age, 1) | ~1 | ~1 | ~1 |
| | PD_PeriodicityWang_th0_01 | -SEP4 | y ~ 1 + fp(age, 1) | ~1 | ~1 | ~1 |
| | SB_BinaryStats_diff_longstretch0 | PE | y ~ 1 + fp(age, 1) | ~1 | ~1 | ~1 |
| | SB_BinaryStats_mean_longstretch1 | PE | y ~ 1 + fp(age, 1) | ~1 | ~1 | ~1 |
| | SB_MotifThree_quantile_hh | ST3 | y ~ 1 + fp(age, 1) | ~1 | ~1 | ~1 |
| | SB_TransitionMatrix_3ac_sumdiagcov | BCT | y ~ 1 + fp(age, 1) | ~1 | ~1 | ~1 |
| | SC_FluctAnal_2_dfa_50_1_2_logi_prop_r1 | ST3 | y ~ 1 + fp(age, 1) | ~1 | ~1 | ~1 |
| | SC_FluctAnal_2_rsrangefit_50_1_logi_prop_r1 | GG | y ~ 1 + fp(age, 1) | ~1 | ~1 | ~1 |
| | SP_Summaries_welch_rect_area_5_1 | -SEP2 | y ~ 1 + fp(age, 1) | ~1 + fp(age, 1) | ~1 | ~1 |
| | SP_Summaries_welch_rect_centroid | -SEP2 | y ~ 1 + fp(age, 1) | ~1 + fp(age, 1) | ~1 | ~1 |
| **Complexity** | app_ent | ST3 | y ~ 1 + fp(age, 1) | ~1 | ~1 | ~1 |
| | detrend_fluc.Alpha | GG | y ~ 1 + fp(age, 1) | ~1 | ~1 | ~1 |
| | detrend_fluc.Beta | ST3 | y ~ 1 + fp(age, 1) | ~1 | ~1 | ~1 |
| | detrend_fluc.Broadband | GG | y ~ 1 + fp(age, 1) | ~1 | ~1 | ~1 |
| | detrend_fluc.Delta | PE | y ~ 1 + fp(age, 1) | ~1 | ~1 | ~1 |
| | detrend_fluc.Gamma | -SEP3 | y ~ 1 + fp(age, 1) | ~1 | ~1 | ~1 |
| | detrend_fluc.Theta | GB2 | y ~ 1 + fp(age, 1) | ~1 | ~1 | ~1 |
| | higuchi_frac.Alpha | GG | y ~ 1 + fp(age, 1) | ~1 | ~1 | ~1 |
| | higuchi_frac.Beta | ST3 | y ~ 1 + fp(age, 1) | ~1 | ~1 | ~1 |
| | higuchi_frac.Broadband | -SEP4 | y ~ 1 + fp(age, 1) | ~1 | ~1 | ~1 |
| | higuchi_frac.Delta | PE | y ~ 1 + fp(age, 1) | ~1 | ~1 | ~1 |
| | higuchi_frac.Gamma | -SEP3 | y ~ 1 + fp(age, 1) | ~1 | ~1 | ~1 |
| | higuchi_frac.Theta | SHASH | y ~ 1 + fp(age, 1) | ~1 | ~1 | ~1 |
| | hjorth_activity | BCT | y ~ 1 + fp(age, 1) | ~1 | ~1 | ~1 |
| | hjorth_complexity | GG | y ~ 1 + fp(age, 1) | ~1 | ~1 | ~1 |
| | hjorth_mobility | ST3 | y ~ 1 + fp(age, 1) | ~1 | ~1 | ~1 |
| | hurst_exp_c | GG | y ~ 1 + fp(age, 1) | ~1 | ~1 | ~1 |
| | hurst_exp_h | exGAUS | y ~ 1 + fp(age, 1) | ~1 | ~1 | ~1 |
| | katz_frac.Alpha | PE | y ~ 1 + fp(age, 1) | ~1 + fp(age, 1) | ~1 + fp(age, 1) | ~1 |
| | katz_frac.Beta | GG | y ~ 1 + fp(age, 1) | ~1 | ~1 | ~1 |
| | katz_frac.Broadband | GG | y ~ 1 + fp(age, 1) | ~1 | ~1 | ~1 |
| | katz_frac.Delta | ST3 | y ~ 1 + fp(age, 1) | ~1 | ~1 | ~1 |
| | katz_frac.Gamma | -SEP4 | y ~ 1 + fp(age, 1) | ~1 + fp(age, 1) | ~1 + fp(age, 1) | ~1 |
| | katz_frac.Theta | GG | y ~ 1 + fp(age, 1) | ~1 | ~1 | ~1 |
| | samp_ent | ST3 | y ~ 1 + fp(age, 1) | ~1 | ~1 | ~1 |
| | spect_ent | exGAUS | y ~ 1 + fp(age, 1) | ~1 | ~1 | ~1 |
| **Spectral** | adjusted_power.Alpha | exGAUS | y ~ 1 + fp(age, 1) | ~1 | ~1 | ~1 |
| | adjusted_power.Beta | exGAUS | y ~ 1 + fp(age, 1) | ~1 | ~1 | ~1 |
| | adjusted_power.Delta | ST3 | y ~ 1 + fp(age, 1) | ~1 | ~1 | ~1 |
| | adjusted_power.Gamma | exGAUS | y ~ 1 + fp(age, 1) | ~1 | ~1 | ~1 |
| | adjusted_power.Theta | GG | y ~ 1 + fp(age, 1) | ~1 | ~1 | ~1 |
| | aperiodic_intercept | ST3 | y ~ 1 + fp(age, 1) | ~1 | ~1 | ~1 |
| | aperiodic_slope | -SEP3 | y ~ 1 + fp(age, 1) | ~1 | ~1 | ~1 |
| | delta_alpha_ratio | ST3 | y ~ 1 + fp(age, 1) | ~1 | ~1 | ~1 |
| | delta_beta_ratio | -SEP2 | y ~ 1 + fp(age, 1) | ~1 + fp(age, 1) | ~1 | ~1 |
| | peak_alpha_freq | GG | y ~ 1 + fp(age, 1) | ~1 | ~1 | ~1 |
| | power_abs.Alpha | GG | y ~ 1 + fp(age, 1) | ~1 | ~1 | ~1 |
| | power_abs.Beta | exGAUS | y ~ 1 + fp(age, 1) | ~1 | ~1 | ~1 |
| | power_abs.Delta | -SEP4 | y ~ 1 + fp(age, 1) | ~1 | ~1 | ~1 |
| | power_abs.Gamma | -SEP2 | y ~ 1 + fp(age, 1) | ~1 | ~1 | ~1 |
| | power_abs.Theta | BCT | y ~ 1 + fp(age, 1) | ~1 | ~1 | ~1 |
| | power_rel.Alpha | exGAUS | y ~ 1 + fp(age, 1) | ~1 | ~1 | ~1 |
| | power_rel.Beta | GG | y ~ 1 + fp(age, 1) | ~1 | ~1 | ~1 |
| | power_rel.Delta | PE | y ~ 1 + fp(age, 1) | ~1 | ~1 | ~1 |
| | power_rel.Gamma | -SEP2 | y ~ 1 + fp(age, 1) | ~1 + fp(age, 1) | ~1 | ~1 |
| | power_rel.Theta | GG | y ~ 1 + fp(age, 1) | ~1 | ~1 | ~1 |
| | theta_alpha_ratio | exGAUS | y ~ 1 + fp(age, 1) | ~1 | ~1 | ~1 |
| | theta_beta_ratio | -SEP2 | y ~ 1 + fp(age, 1) | ~1 + fp(age, 1) | ~1 | ~1 |
| **Time** | env_mean.Alpha | SHASHo | y ~ 1 + fp(age, 1) | ~1 | ~1 | ~1 |
| | env_mean.Beta | GG | y ~ 1 + fp(age, 1) | ~1 | ~1 | ~1 |
| | env_mean.Broadband | BCT | y ~ 1 + fp(age, 1) | ~1 | ~1 | ~1 |
| | env_mean.Delta | BCT | y ~ 1 + fp(age, 1) | ~1 | ~1 | ~1 |
| | env_mean.Gamma | ST3 | y ~ 1 + fp(age, 1) | ~1 | ~1 | ~1 |
| | env_mean.Theta | PE | y ~ 1 + fp(age, 1) | ~1 | ~1 | ~1 |
| | env_std.Alpha | SHASHo | y ~ 1 + fp(age, 1) | ~1 | ~1 | ~1 |
| | env_std.Beta | GG | y ~ 1 + fp(age, 1) | ~1 | ~1 | ~1 |
| | env_std.Broadband | SHASHo | y ~ 1 + fp(age, 1) | ~1 | ~1 | ~1 |
| | env_std.Delta | -SEP2 | y ~ 1 + fp(age, 1) | ~1 | ~1 | ~1 |
| | env_std.Gamma | ST3 | y ~ 1 + fp(age, 1) | ~1 | ~1 | ~1 |
| | env_std.Theta | PE | y ~ 1 + fp(age, 1) | ~1 | ~1 | ~1 |
| | kurt_t.Alpha | ST3 | y ~ 1 + fp(age, 1) | ~1 + fp(age, 1) | ~1 | ~1 |
| | kurt_t.Beta | JSU | y ~ 1 + fp(age, 1) | ~1 | ~1 | ~1 |
| | kurt_t.Broadband | exGAUS | y ~ 1 + fp(age, 1) | ~1 + fp(age, 1) | ~1 | ~1 |
| | kurt_t.Delta | exGAUS | y ~ 1 + fp(age, 1) | ~1 | ~1 | ~1 |
| | kurt_t.Gamma | SHASHo | y ~ fp(age, 1) + factor(sex) | ~fp(age, 1) + factor(sex) | ~factor(sex) | ~1 |
| | kurt_t.Theta | exGAUS | y ~ 1 + fp(age, 1) | ~1 | ~1 | ~1 |
| | power_t.Alpha | GB2 | y ~ 1 + fp(age, 1) | ~1 | ~1 | ~1 |
| | power_t.Beta | exGAUS | y ~ 1 + fp(age, 1) | ~1 | ~1 | ~1 |
| | power_t.Broadband | BCT | y ~ 1 + fp(age, 1) | ~1 | ~1 | ~1 |
| | power_t.Delta | PE | y ~ 1 + fp(age, 1) | ~1 | ~1 | ~1 |
| | power_t.Gamma | -SEP2 | y ~ 1 + fp(age, 1) | ~1 | ~1 | ~1 |
| | power_t.Theta | BCT | y ~ 1 + fp(age, 1) | ~1 | ~1 | ~1 |
| | skew_t.Alpha | JSU | y ~ 1 + fp(age, 1) | ~1 | ~1 | ~1 |
| | skew_t.Beta | ST3 | y ~ 1 + fp(age, 1) | ~1 | ~1 | ~1 |
| | skew_t.Broadband | -SEP2 | y ~ 1 + fp(age, 1) | ~1 | ~1 | ~1 |
| | skew_t.Delta | ST3 | y ~ 1 + fp(age, 1) | ~1 | ~1 | ~1 |
| | skew_t.Gamma | SHASH | y ~ 1 + fp(age, 1) | ~1 | ~1 | ~1 |
| | skew_t.Theta | -SEP4 | y ~ 1 + fp(age, 1) | ~1 | ~1 | ~1 |
